# cLD: Rare-variant disequilibrium between genomic regions identifies novel genomic interactions

**DOI:** 10.1101/2022.02.16.480745

**Authors:** Dinghao Wang, Jingni He, Deshan Perera, Chen Cao, Pathum Kossinna, Qing Li, William Zhang, Xingyi Guo, Alexander Platt, Jingjing Wu, Qingrun Zhang

**Author notes:** joint first authors (Dinghao Wang, Jingni He, Deshan Perera). Correspondence should be address to Q.Z.

## Abstract

Linkage disequilibrium (LD) is a fundamental concept in genetics; critical for studying genetic associations and molecular evolution. However, LD measurements are only reliable for common genetic variants, leaving low-frequency variants unanalyzed. In this work, we introduce cumulative LD (cLD), a stable statistic that captures the rare-variant LD between genetic regions, which reflects more biological interactions between variants, in addition to lack of recombination. We derived the theoretical variance of cLD using delta methods to demonstrate its higher stability than LD for rare variants. This property is also verified by bootstrapped simulations using real data. In application, we find cLD reveals an increased genetic association between genes in 3D chromatin interactions, a phenomenon recently reported negatively by calculating standard LD between common variants. Additionally, we show that cLD is higher between gene pairs reported in interaction databases, identifies unreported protein-protein interactions, and reveals interacting genes distinguishing case/control samples in association studies.

## INTRODUCTION

Linkage Disequilibrium (LD) is a fundamental concept in population genetics that statistically captures non-random associations between two genetic variants due to reasons such as lack of recombination or different age of mutations (Slatkin 2008). LD serves as a core component in genotype-phenotype association mapping, as a statistically significant genetic variant could be just a proxy in LD with the genuine causal variant(s) (Weissbrod et al. 2020). To this end, LD is critically important in analyzing the fine resolution of genotype-phenotype association mapping (Flint-Garcia et al. 2003) and forming polygenic risk scores (Amariuta et al. 2020). Additionally, from the perspective of molecular evolution, LD values substantially higher than expected under neutrality may indicate interesting phenomena, e.g., interactions between loci that are favored by selection (Gregersen et al. 2006). As such, LD has been extensively utilized in evolutionary studies.

The calculation of LD involves the use of allele frequencies of the genetic variants in its denominator to normalize the statistic (**Methods**; **Supplementary Materials 1.1**) and therefore suffers from a high variance (instability) when allele frequencies are close to zero. As such, in practice, researchers only analyze common genetic variants with minor allele frequency (MAF) higher than a threshold (e.g., 0.05), excluding more than 90% of human genetic variants (Auton et al. 2015).

In the field of association mapping, researchers have developed multiple techniques to aggregate the associations of multiple rare variants with a phenotype into a single shared effect. One of the pioneering methods that is still popularly used (Li and Leal 2008) is synthesizing a cumulative allele frequency from multiple rare genetic variants in the same genetic region (e.g., within a gene). The cumulative minor allele frequency (cMAF) is defined on a region containing multiple rare variants: an individual will be labelled as a “mutant” if it has at least one of the rare variants, and then the proportion of individuals in the sample that are labelled as mutants will be the cMAF for this region (**Fig. 1a**).

**Figure 1.**
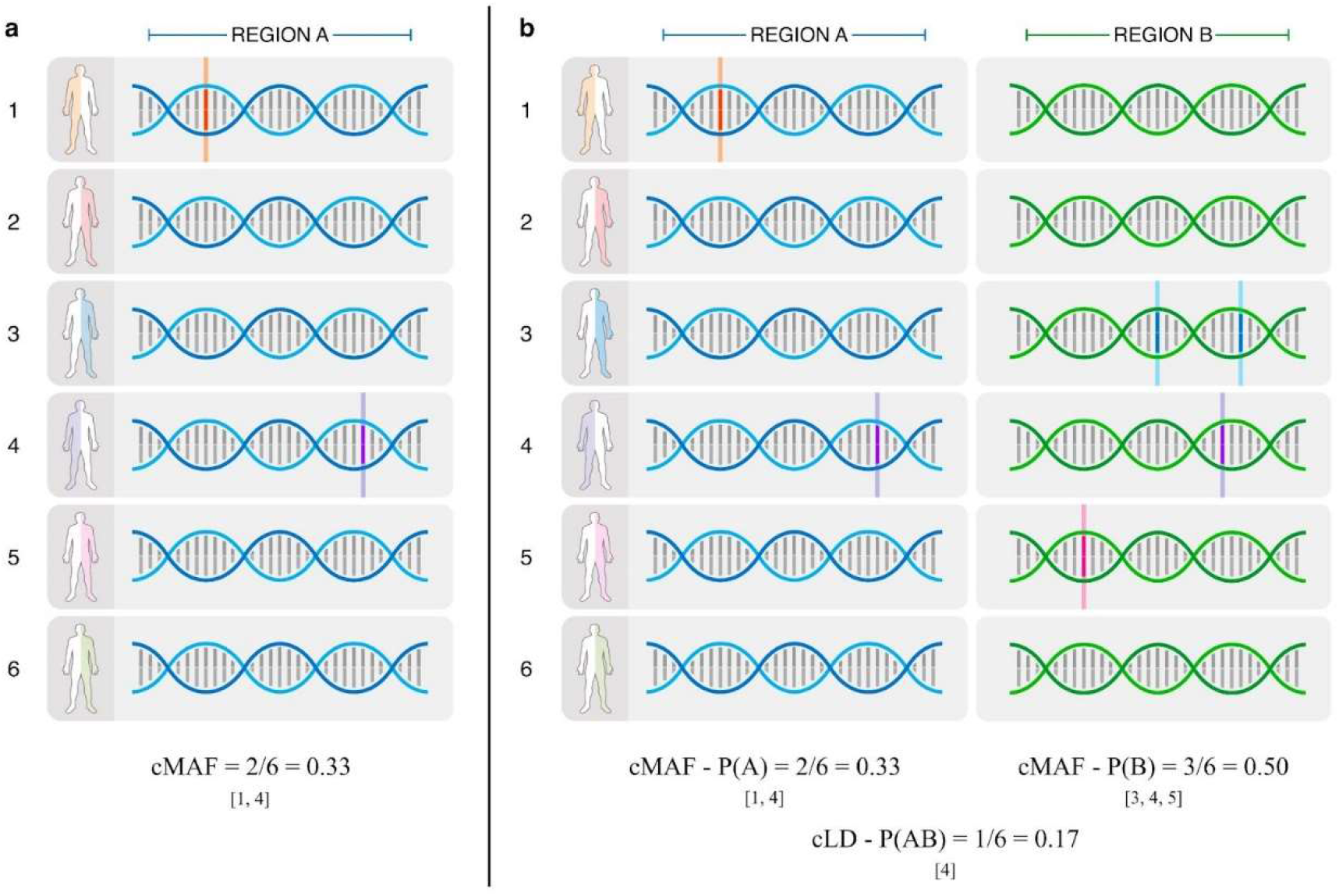
Illustration of the idea of a) cMAF and b) cLD. An example to show the calculation of cLD, inspired by cMAF. **a)** Out of six haplotypes, there are two [1, 4] who have mutations in region A. Therefore, the cMAF P(A) for region A is 2/6 = 0.33. **b)** There are three haplotypes [3, 4, 5] who have mutations in region B and the cMAF P(B) for region B is 3/6 = 0.50. If one considers regions A and B together, there is one individual with mutations in both regions: [4]. Thus, the P(AB) is 1/6 = 0.17. Finally, by yielding P(A), P(B) and P(AB) into the standard formula of LD we have cLD = 0.375.

Building on the idea of cMAF and the essence of LD, we developed a statistic, cumulative Linkage Disequilibrium (cLD) to capture the aggregated correlation between two sets of rare variants (**Methods; Fig. 1b**).

We thoroughly tested the property of cLD. First, using both theoretical closed-form derivation and bootstrapped simulations (**Methods**), it is verified that cLD enjoys way lower variance than the standard LD when applied to rare variants, evidencing cLD’s higher stability (**Fig. 2**). We then applied cLD to four scenarios in genetic analysis (**Methods**), discovering additional knowledge that have not been reported (or attempted but negatively reported) using standard LD (**Figs. 3 – 6**).

**Figure 2.**
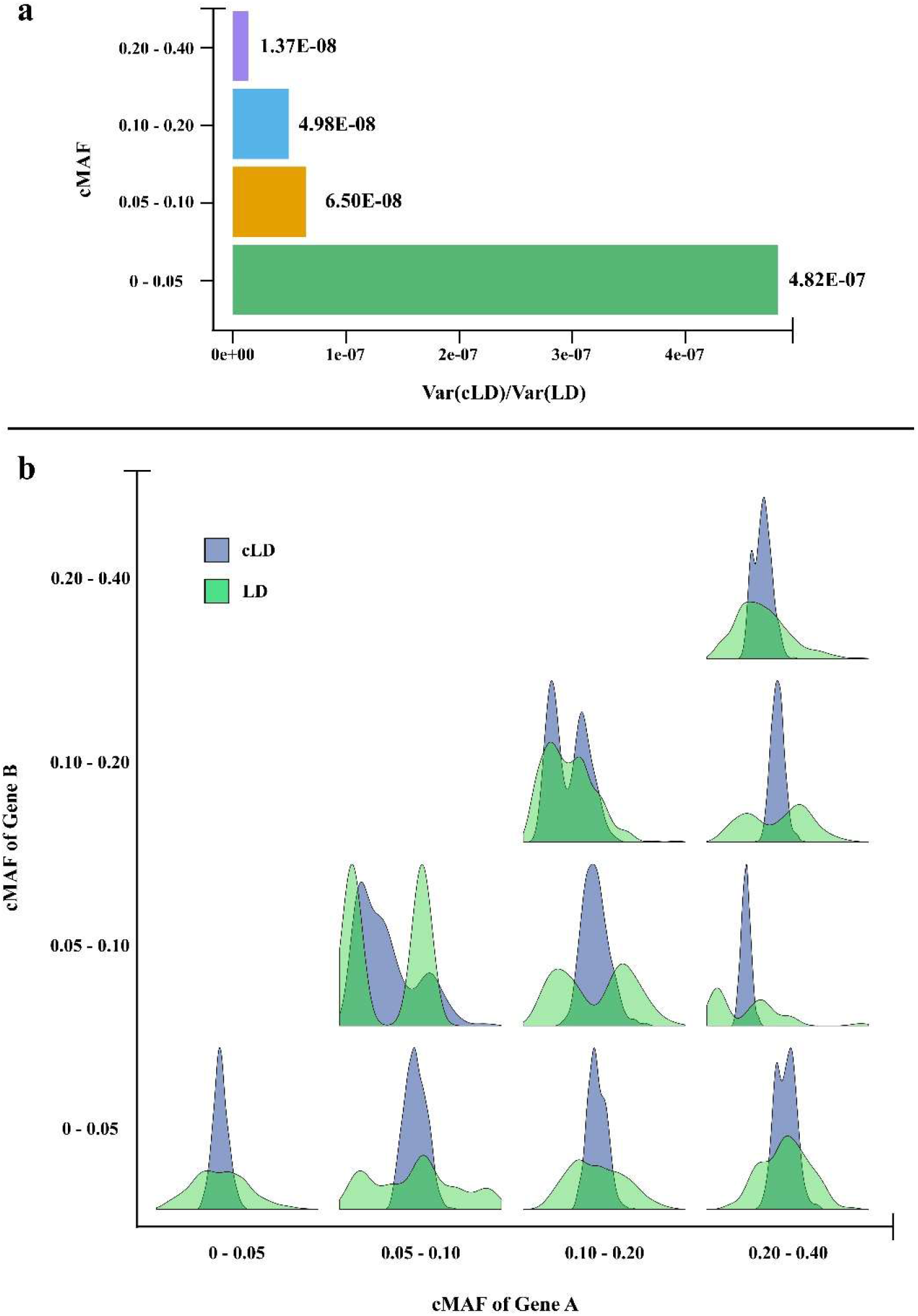
Stability of cLD and LD revealed by closed-form variance calculation and bootstrapped distributions. **a)** The gene pairs were split into four different bins based on the cMAF values, i.e., <0.05, 0.05 - 0.10, 0.10 - 0.20, and 0.20 - 0.40 (y-axis). The x-axis is the ratio between the variances of cLD and LD, i.e., Var(cLD)/Var(LD). **b)** Probability density distribution of cLD and LD from bootstrapped samples. Results from the European population are shown. See **Supplementary Figs. <u>S</u>2.1 & <u>S</u>2.2** for other populations.

**Figure 3.**
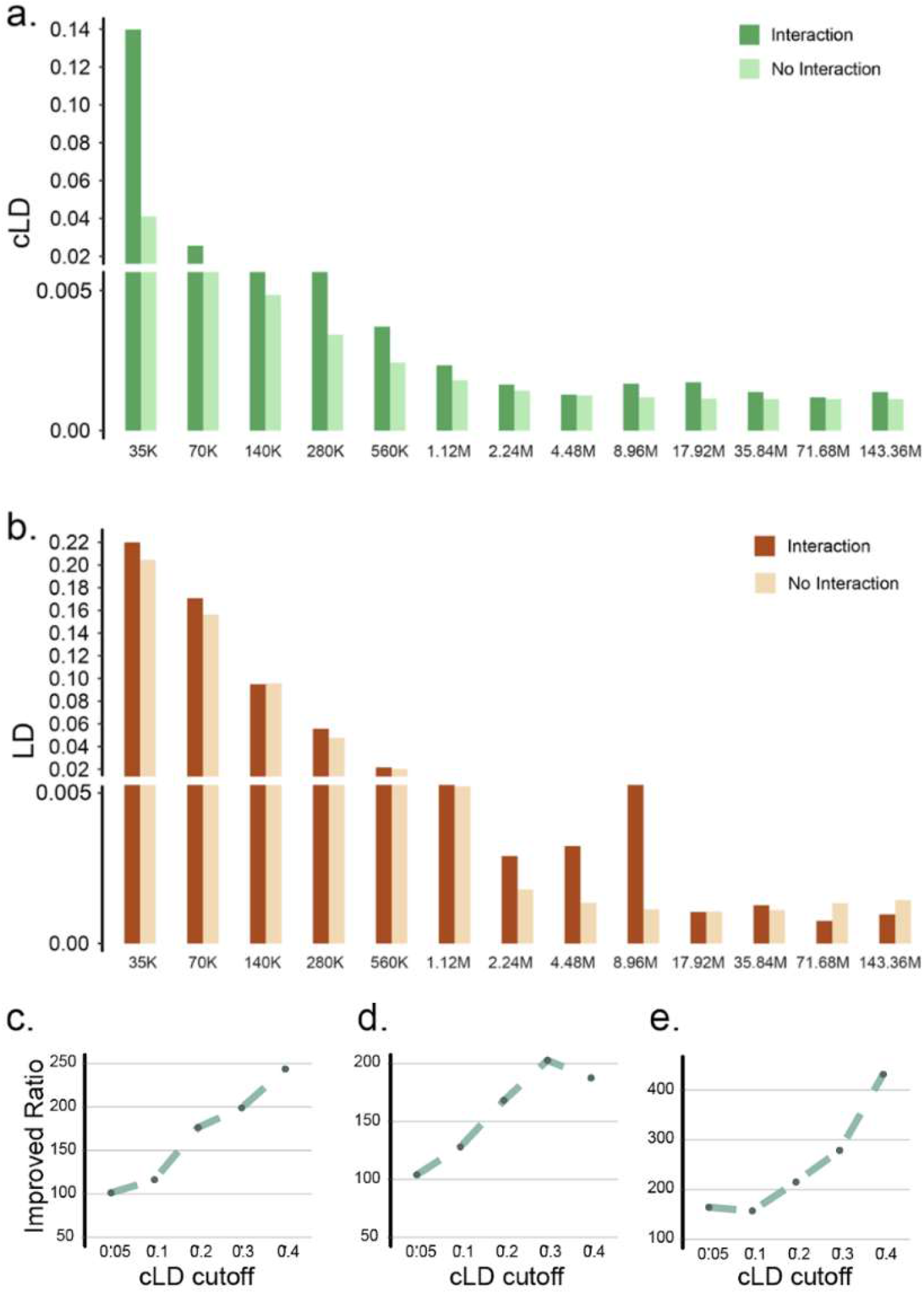
Enrichment of cLD among pairs of genes in chromatin contact regions. **a)** The comparisons of cLD values between the 3D chromatin interaction regions and non-interaction regions among 13 different distance groups in the European population. (Other populations are shown in **Supplementary Fig. <u>S</u>3.1**) The confidence intervals for these bars are presented in **Supplementary Table <u>S</u>3.1**. **b)** The same comparisons using standard LD in the European population. (Other populations are shown in **Supplementary Fig. <u>S</u>3.2**) **c-e)** The ratios between the number of gene pairs in 3D chromatin interaction regions against the number of gene pairs that are not in 3D regions. The x-axis is the cLD value cutoffs above which the gene pairs are counted. **c)** European population. **d)** African population. **e)** East Asian population.

**Figure 4.**
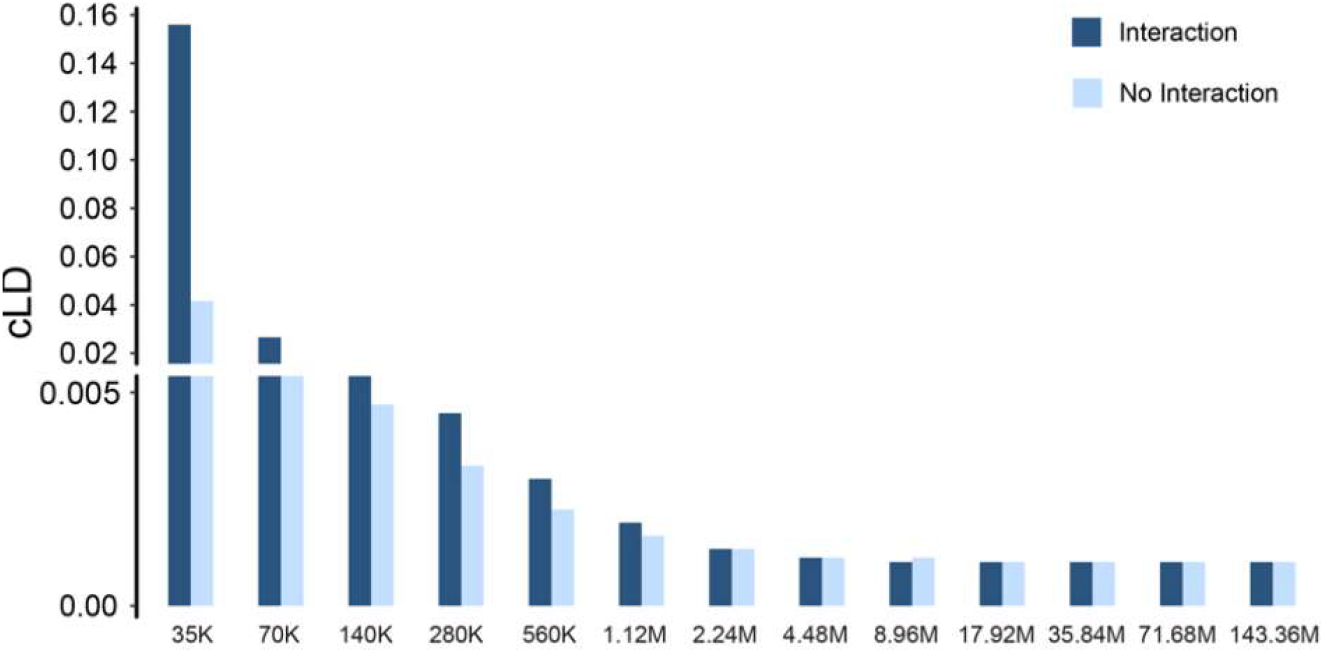
The comparisons of cLD values in European populations between gene pairs found in interaction databases and all pairs that are not in databases. Each bar represents the average of pairs with distance smaller than the value of its x-axis label but larger than the value of the previous x-axis label. (Other populations show the same trend, as depicted in **Supplementary Fig. <u>S</u>3.3**)

**Figure 5:**
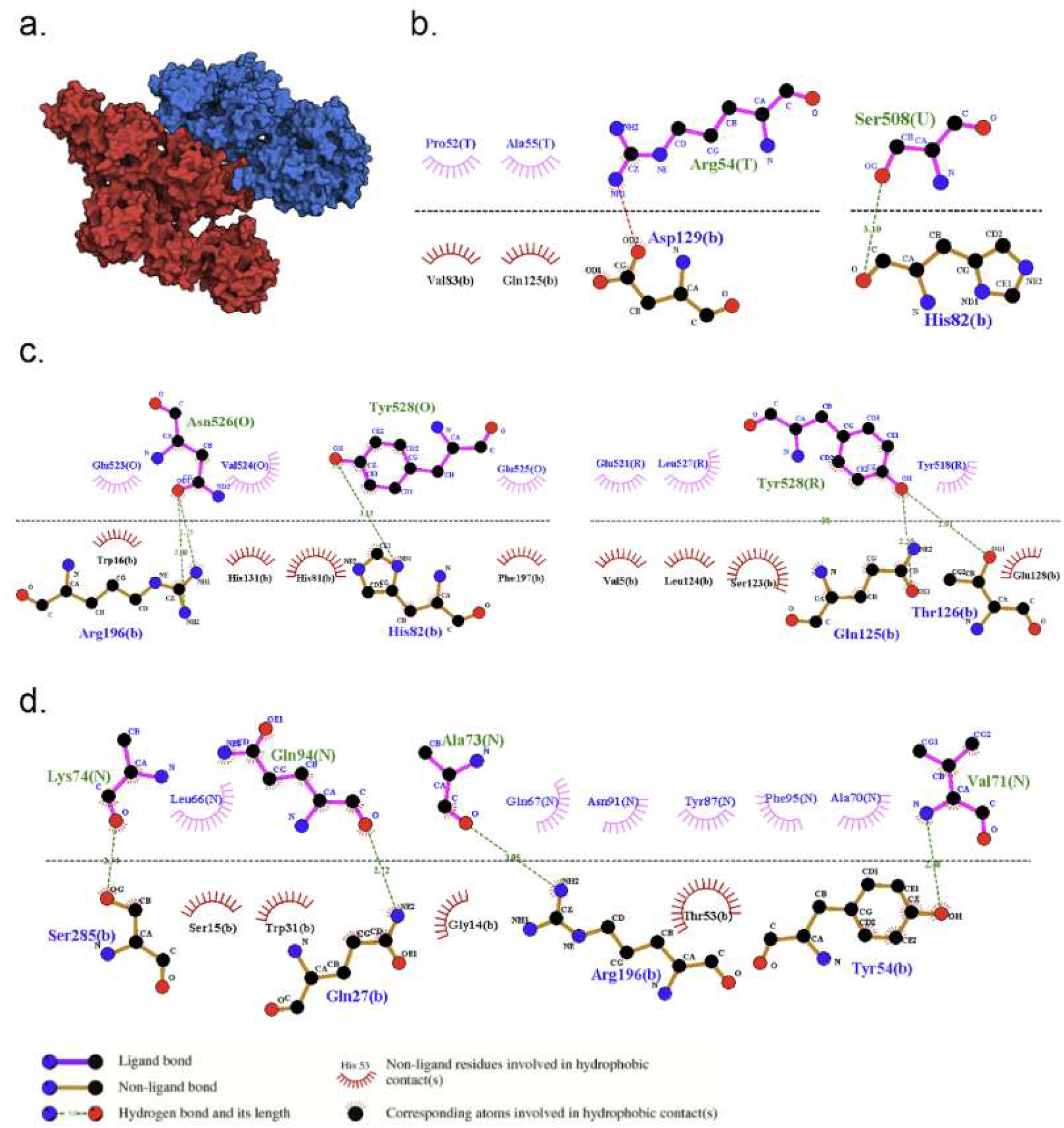
Protein docking interaction between 3BCZ and 4RIQ revealed by cLD (=0.86) with a binding affinity of −341.21 kJ/mol. **a)** Structure of 3BCZ (red) and 4RIO (blue) protein-protein complex. **b-d)** 2D representation of closest interacting residues around the protein-protein interaction interfaces, including multiple non-covalent bonds, for example, hydrogen bonds (green dotted line) and hydrophobic interactions (read and rose semi-circle with spikes). Residues for the 3BCZ are depicted in upper letters (T, U, O, R, N) and for the 4RIO are depicted in lower letters.

**Figure 6:**
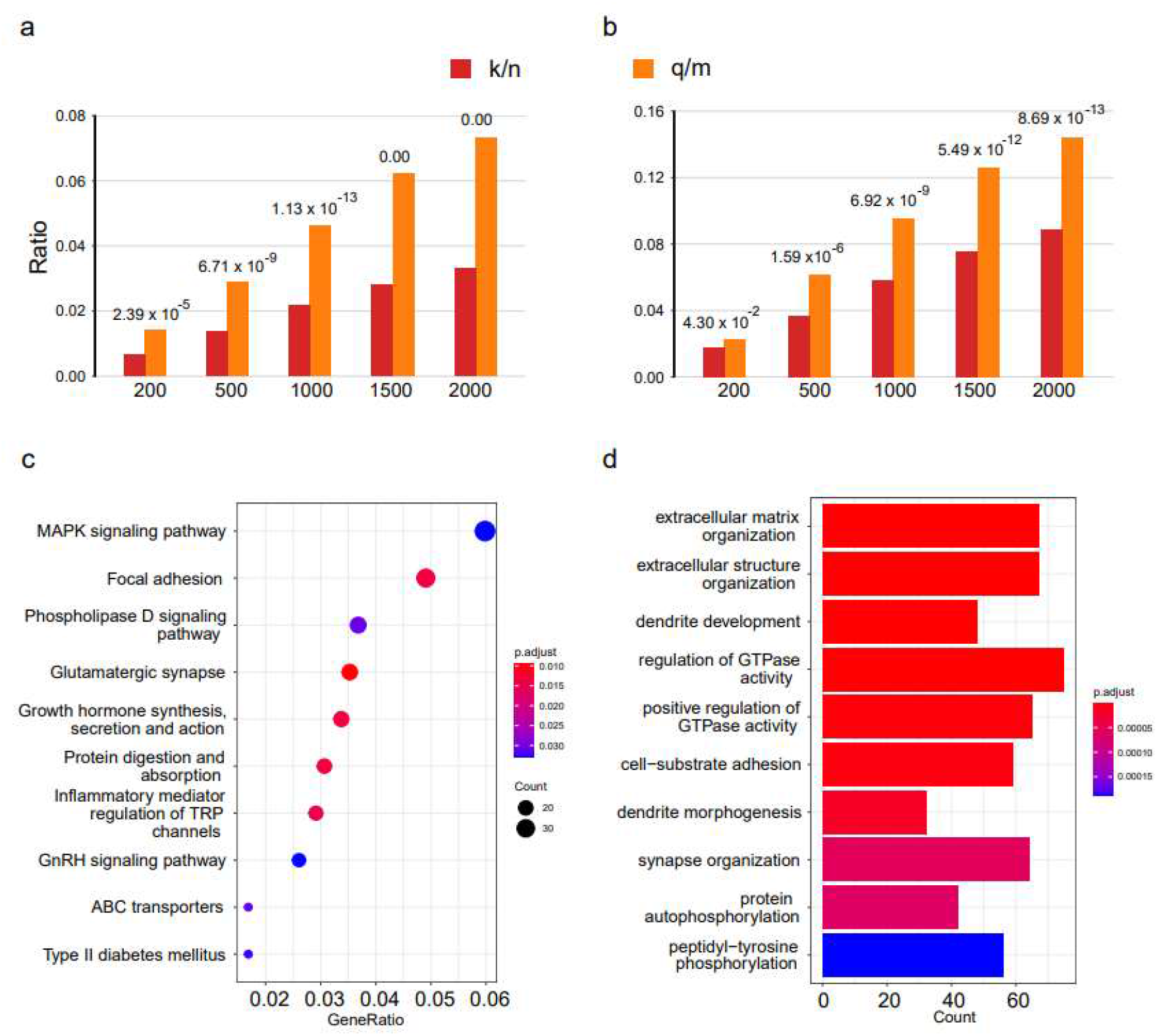
ΔcLD gene pairs in case/control association mapping data: annotation of top genes and enrichment of pathways. **a-b)** Group bar charts show the ratio between the number of selected genes being validated in the database dividing the number of genes in the database (q/m) as well as the number of selected genes dividing the total number of all known minus m (k/n). The values on the top of each bar are the p-values of the hypergeometric distribution probability test. The x-axis indicated the top gene pairs using different cutoffs, [200, 500, … 2,000]. **a)** DisGeNET database. **b)** SFARI database. **c)** a dot plot showing the top 10 KEGG pathways ranked by the GeneRatio values. The size of the balls indicates the number of the genes enriched and the color indicates the level of the enrichment (P-adjusted values). The GeneRatio is calculated as count/setSize. ‘count’ is the number of genes that belong to a given gene-set, while ‘setSize’ is the total number of genes in the gene-set. **d)**. a bar plot showing the top 10 enriched biological processes ranked by p-values. The correlation is more significant as the red/blue ratio increases. The number on the x-axis indicates the number of genes that belong to a given gene set.

## RESULTS

### The intuitive idea of defining cLD

In the similar vein of definition of cMAF, we define cLD below. Specifically, for the traditional calculation of LD between two variants, *g1* and *g2* with minor alleles *a* and *b* respectively, the essential part is the definition of individual MAF *P*(*a*) and *P*(*b*) and the frequency that a and b show up in the same haplotype, *P*(*ab*). For calculating cLD between two regions, *A* and *B*, we first use cMAF to define *P*(*A*) and *P*(*B*) (the proportion of individuals carrying a rare variant within regions *A* and B, respectively); and then *P*(*AB*), the proportion of individuals who have at least one rare variant in both regions A and B (**Fig. 1b**). Mathematical details are spelt out in **Methods** and **Supplementary Materials 1.1 & 1.2**).

### High stability of cLD in contrast to standard LD

Both LD and cLD could be used to capture the correlation between two sets of rare variants. However, these two measures differ in the aspect of stability. Intuitively, as cMAF is always higher than MAF, cLD’s variance (reflecting its instability) should be lower than LD’s. We verify this intuition by deriving the closed-form of variance of both LD and cLD (denoted as Var(LD) and Var(cLD)) using multinomial distributions and their multivariate normal approximation as well as the multivariate Delta Method (Lehmann Springer) (**Methods; Supplementary Materials 2.1 & 2.2**). by plugging in the allele frequencies calculated using the 1000 Genomes Project data (Auton et al. 2015) (**Supplementary Materials 2.3**), we observed that the variance of cLD is at least six orders of magnitudes smaller (i.e., more stable) than the alternative -- calculating LD directly on rare variants in all ethnic populations and all cMAF bins (**Fig. 2a; Supplementary Figs. S2.1a & S2.2a**). Additionally, following the conventional statistical procedure of bootstrapping to empirically estimate stability, we sub-sampled half of each population sample 1,000 times to form bootstrapped distributions for both cLD and LD (**Methods; Supplementary Materials 2.4**). The subsampling showed that cLD exhibits much slimmer bootstrapped distributions than LD across all cMAF bins and all three ethnic groups (**Fig. 2b, Supplementary Figs. S2.1b & S2.2b**), further confirming the greater stability of cLD compared to traditional measures of LD.

### cLD reveals linkage disequilibrium between 3D contact regions where standard LD fails

A distinct advantage of cLD over LD is the ability to reveal linkage disequilibrium between 3D contact regions. By aggregating information from multiple independent mutations, cLD is sensitive to subtle interactions poorly reflected by LD (which can only account for two at a time). As such, cLD captures more biological interactions in addition to traditional LD that focuses more on the lack of recombination. Interactions within the 3D structure of genomes is one place where this difference allows for insight from cLD where LD-based methods fail. The availability of high-throughput experimental technologies that can assess chromatin conformation such as Hi-C (Rajarajan et al. 2018; Akbarian et al. 2015) allows researchers to analyze genetic regions that are in close contact in 3D spatial structure. There was a widely disseminated expectation that the 3D genomic interaction in the form of chromatin contact may leave a footprint in the form of genetic LD (Joiret et al. 2019). Motivated by such expectation, Whalen and Pollard calculated the standard LD based on common variants (MAF>0.05) in 1000 Genomes Project data (Auton et al. 2015) and reported negative results stating that genetic LD map is not overlapping with the 3D contact map (Whalen and Pollard 2019). However, by reanalyzing the 1000 Genomes sequencing data and Hi-C data (Akbarian et al. 2015; Rajarajan et al. 2018) in the developing brain using cLD on rare variants (**Methods; Supplementary Materials 3.1 & 3.2**), we revealed that the 3D chromatin interactions did leave genetic footprints in the form of higher cLD in pairs of genes that are in the adjacent Hi-C regions (**Fig 3a; Supplementary Fig. S3.1**). To assess the statistical significance of the enrichment of cLD in 3D contact regions, we conducted Mantel-Haenszel and Fisher exact tests (**Supplementary Materials 3.4**), both of which are highly significant (P-value < 1.0E-50; **Supplementary Tables S3.2 & S3.6, Supplementary Materials 3.4.1**). As Whalen & Pollard’s work (Whalen and Pollard 2019) is not at the resolution of genes, we re-calculated standard LD using common variants based on gene pairs (**Supplementary Materials 3.2**), which shows a subtle effect (**Fig. 3b, Supplementary Fig. S3.2**) but still not statistically significant with Mantel-Haenszel and Fisher exact tests (P-value =0.999; **Supplementary Tables S3.3 & S3.4; Supplementary Materials 3.4.1**). Additionally, we checked the ratio between the number of pairs of genes within the 3D contact regions and the number of pairs outside the 3D contact regions as a function of their cLD cut-off. More specifically, we prespecified a cLD value cutoff and only counted the gene pairs with cLD value higher than this cutoff; then we separated the number of genes within or outside 3D contact regions and calculated their ratios (**Supplementary Materials 3.5**). Indeed, we found that the ratios are significantly larger than 1.0 and increase as the cLD cutoffs increase (**Fig 3c,d,e, Supplementary Table S3.7)**. Taking together, 3D interactions clearly overlap with genetic interactions; and cLD is instrumental in observing this whereas standard LD fails.

### cLD is enriched in known interacting genes

To demonstrate that gene-gene interactions leave footprints in rare genetic mutations regardless of their physical positions we computed the distribution of cLD among interacting pairs genes reported in Reactome (Fabregat et al. 2018) and BioGRID (Stark et al. 2006), MINT (Orchard 2012) and IntAct (Orchard et al. 2014) (**Methods; Supplementary Materials 3.3**). We compared this distribution against a null distribution formed by all pairs of genes. Indeed, the comparisons led to the expected result: for gene pairs separated by any physical distance within 2MB, cLD is elevated in interacting gene pairs (**Fig. 4; Supplementary Fig. S3.3**). Again, the Mantel-Haenszel and Fisher exact tests confirm that the differences are significant (P-value < 1.0E-20; **Supplementary Table S3.5; Supplementary Materials 3.4.2**).

### cLD identified novel pairs of likely interacting proteins

To examine the novel gene pairs with higher cLD values have the receptor-ligand interactions of their translated proteins, we performed protein-docking analysis to obtain the evidence. Looking at all pairs of genes, we observed several pairs without prior evidence of interaction with extraordinarily high cLD, such as between genes *MEMO1* and *DPY30* (encoding proteins 3BCZ and 4RIQ, respectively) with a cLD of 0.86. We conducted protein docking analysis for the genes of large cLD values (top 0.01% among all gene pairs) with cMAF > 0.05 and existing IDs in PDB, however, not reported in any interaction databases (**Methods; Supplementary Materials 4.1; Supplementary Table S4.1**). These criteria lead to 19 pairs of genes for protein-docking. We found multiple lines of evidence of the interaction at protein level for five pairs (**Supplementary Table S4.2**) in terms of both binding affinity and interacting residues (**Fig. 5a-d; Supplementary Figs. S4.1 - S4.4**).

### Differences in cLD distinguish cases/controls in Autism exome data

In the context of case/control association studies, cLD can be used to identify pairs of genes whose interactions may be responsible for human diseases. Using data from the *Autism Spectrum Disorders* (ASD) whole exome sequencing dataset (Satterstrom et al. 2020), we calculated cLD values for all pairs of genes, separately conducted for the populations of cases and controls (**Methods; Supplementary Materials 5.1 & 5.2**). The difference in cLD for a pair of genes conditional on case/control status, defined as ***ΔcLD***, is indicative of an interaction that is non-random associating with disease status. We collected gene pairs with high ΔcLD and checked their annotation and enrichment in existing databases. Using a hypergeometric test, we analyzed the enrichment among high-ΔcLD genes for ASD genes reported by DisGeNet (Piñero et al. 2017), an established general database for diseases and SFARI (Abrahams et al. 2013), a gold-standard database focusing on ASD (**Supplementary Materials 5.3**). The genes included in the pairs with high ΔcLD scores are highly enriched in both the Autism related genes in DisGeNet (**Fig. 6a**) and SFARI (**Fig. 6b**). Gene Ontology (Ashburner et al. 2000) and pathways (KEGG) (Kanehisa and Goto 2000; Kanehisa et al. 2009) enrichment analysis for the high ΔcLD genes (**Methods; Supplementary Table S5.2; Supplementary Materials 5.4**) also showed sensible biological functions and pathways (**Fig. 6c,d**) that are well supported by the literature (**Supplementary Materials 5.4**) (Ashburner et al. 2000; Kanehisa and Goto 2000; Kanehisa et al. 2009; Yu et al. 2012; Rojas 2014; Hannelius et al. 2005; Richler et al. 2006; O’Roak et al. 2012; Fung and Hardan 2015; Sato et al. 2012; Berkel et al. 2010; Durand et al. 2007; Wei et al. 2021; Ye et al. 2011; Betancur et al. 2009; Lin et al. 2016). By taking a closer look of the 20 genes identified by the top 10 gene pairs with the highest ΔcLD values, found that 14 genes (70%) have been reported to be associated with ASD, including *DENND4A, EFCAB5, ABI2, RAPH1, MSTO1, DAP3, ARL13B, PRB2, PRB1, ZNF276, FANCA, ADAM7, SLC26A1* and *TUBB8* (**Supplementary Table S5.1**). Moreover, among the rest of six genes, we also identified indirect links of two, *RAB11A* and *IDUA* with ASD (**Supplementary Materials 5.3**).

## DISCUSSION

LD is a critical concept applicable to many types of genetic analyses. In this work, we have defined cLD, a new statistic addressing the association between genetic regions using rare genetic variants. In contrast to the previous attempts to utilize LD between multiple variants focusing on dominant haplotypes (Zan et al. 2018) or joint distributions (Turkmen and Lin 2017), cLD emphasizes biological interactions. Additionally, previously researchers have proposed composite linkage (Hamilton and Cole 2004; Zaykin 2004), which addresses the property of variances and its normalization, however, does not incorporate rare variants.

By both closed-form derivations and statistical simulations, we proved the stability of cLD in contrast to the high instability of standard LD (when applied to rare variants). The stability and the focus on biological interaction allows cLD to capture additional information from the distributions of many variants segregating in a population at low frequencies within particular regions of a genome. Indeed, by applying cLD to real data, we observed interesting overlapping pattern of 3D interactions and genetic interactions that have been negatively reported by using standard LD. We also successfully analyzed protein docking and association mapping, providing two broadly impactable use-cases of cLD. With its demonstrated power in identifying gene and protein interactions, cLD might offer an essential tool to analyze biological interactions and their evolution using rare genetic variants.

## METHODS

### Definition of LD and cLD

The definition of LD between two bi-allelic loci relies on the calculation of three key quantities: *P_A_*, the allele frequency of an allele in locus *A, P_B_*, the allele frequency of an allele in locus *B*, and *P_AB_*, the frequency of these two alleles of A and *B* showing up together. Then one can define the unnormalized disequilibrium statistic *D* = *P_AB_* – *P_A_P_B_*. To rescale the statistic based on allele frequency, one can normalize *D* by dividing it by the allele frequency variances:

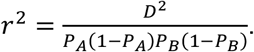

An alternative definition of LD is *D*′, which has a different way of normalization. In this paper, we used *r*^2^ as the representative. Because LD involves *P_A_* and *P_B_* in the denominator, it is highly instable when *P_A_* or *P_B_* are close to zero, which means LD cannot be used if *A* or *B* are rare variants.

The cLD statistic is designed to handle the above problem by aggregating rare variants cumulatively. In the similar vein of definition of cMAF, the idea of cLD is illustrated in **Fig. 1b**. More specifically, here we look at two sets of variants in two genetic regions, e.g., two genes, again namely A and B. Assuming that there are *m* SNPs in gene A, and there are *r* SNPs in gene B. Also, we assume the sample size is *n*. Then, for gene *A*, we use *S*_1*i*_, *S*_1*i*_, …, *S_mi_* to denote the allele of the *s*-th SNP (*s* = 1, 2, …, *m*) in the *i*-th individual (*i* = 1, 2, …, *n*). Similarly, for gene B, we use {*K*_1*i*_, *K*_2*i*_,…, *K_ri_*} to denote the allele of the *k*-th SNP (*k* = 1, 2, …, *r*) in the *i*-th individual (*i* = 1, 2, …, *n*). Note that *S_si_* and *K_ki_* is either 0 or 1. (0 denotes a major allele, whereas 1 denotes a minor allele).

Then we have the cMAF (*P_A_* & *P_B_*) defined below:

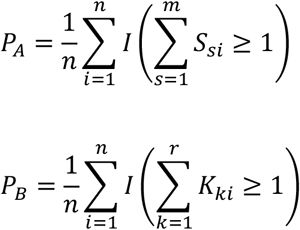

Where *I*(.) is the indicator function. *P_AB_* is then defined as the proportion of individual haplotypes with a minor allele in both regions:

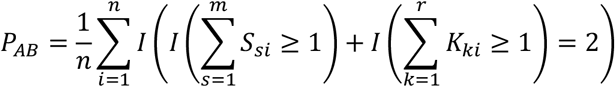

Following the convention of LD, we define the *r*^2^ version of cLD:

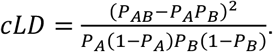

The more rigorous mathematical descriptions and the definition of *D*′ version is provided in **Supplementary Materials 1.1 & 1.2**.

### Derivation of theoretical variance of cLD in contrast to LD

To obtain the theoretical variance of cLD and LD, we derived their asymptotic distributions. The details are in **Supplementary Materials 2.1 & 2.2**. The gist of our approach is summarized in the following three steps:

First, we rewrote the formula of cLD and LD in terms of counts to use multinomial random variables. In the definition, we used *X_ijk_* to denote the allele of the *k*-th variant of the *j*-th gene for the *i*-th individual (haplotype) of. For a pair of variants, the *i*-th pair (*X*_*i*1*u*_, *X*_*i*2*v*_) (*i* = 1, 2, …, *n*) can take possible values (1, 1), (0, 1), (1, 0) and (0, 0). Using *O*_1_ to *O*_4_ to denote the count of the 4 possible pairs in two variants, the distribution of ***O*** = (*O*_1_, *O*_2_, *O*_3_, *O*_4_) is ***O**~multinom*(*n*; ***p***) with ***p*** = (*p*_1_, *p*_2_, *p*_3_, *p*_4_) represents the population probability. The LD between the *u*-th and *v*-th variants can be re-written as:

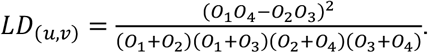

Similarly, we followed the same strategy of using multinomial random variables to describe cLD as below:

In analogy to the case of LD, we used *X*_*ij*_ to denote the allele of the *j-th* gene for the *i-th* individual (haplotype). For a pair of genes, the *i*-th pair (*X*_*i*1_, *X*_*i*2_) (*i* = 1, 2, …, *n*) can take possible values (1, 1), (0, 1), (1, 0) and (0, 0). Using *M*_1_ to *M*_4_ to denote the counts of the 4 possible pairs in two genes, then the distribution of ***M*** = (*M*_1_, *M*_2_, *M*_3_, *M*_4_) is ***M**~multinom*(*n; q*) with ***q*** = (*q*_1_, *q*_2_, *q*_3_, *q*_4_) represents the population probability. The cLD between a pair of genes could be rewritten as:

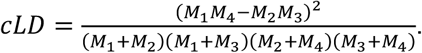

Second, we used the central limit theorem (CLT) to derive the asymptotic multivariate normal distribution. In the LD case, with the population mean ***p*** = (*p*_1_, *p*_2_, *p*_3_, *p*_4_), we can write the covariance matrix as

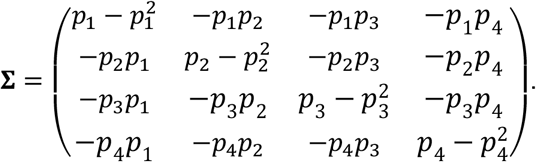

Then by the multivariate CLT (Lehmann Springer) we have 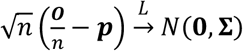.

In the cLD case, with the population mean ***q*** = (*q*_1_, *q*_2_, *q*_3_, *q*_4_), we can write the covariance matrix as

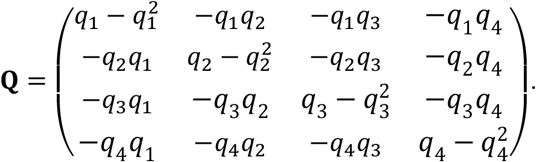

Then by the multivariate CLT (Lehmann Springer) we have 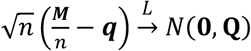.

Third, as the cLD and LD are functions of random variables, we applied the multivariate Delta method (Lehmann Springer) to derive the distribution of cLD and LD. In the LD case, suppose the Jacobian matrix of *LD*(***O***/*n*) is 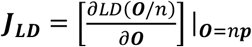. Then the asymptotic distribution of *LD*(***O***/*n*) is 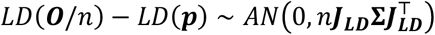, where *‘AN’* stands for asymptotic normal.

In the cLD case, suppose the Jacobian matrix of *cLD*(***M**/n*) is 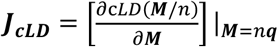. Then the asymptotic distribution of *cLD*(***M**/n*) is 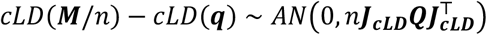.

### Genotype data used for the calculations

The 1000 Genomes Variant Call Data were used to validate the properties of cLD. In particular, the phased (i.e., haploid instead of diploid) variant call data of the Phase 3 of the 1000 Genomes dataset was obtained through The European Bioinformatics Institute’s dedicated FTP server (Fairley et al. 2020).

### Assessing the instability of LD and cLD using bootstrapped distributions

To use bootstrapped samples to quantify instability, we randomly sampled half of the haplotypes in three main 1000 Genomes Project populations (EUR, AFR, or EAS), and calculated the average cLD and average LD over the gene pairs within cMAF bins and repeated this procedure 1,000 times. Based on these bootstrapped cLD and LD values we formed bootstrapped distributions for cLD and LD respectively (with appropriate rescaling described in **Supplementary Materials 2.3**). More specifically, we randomly sampled 1,000 genes and assessed their pairwise LD and cLD in stratified cMAF bins (**Supplementary Materials 2.4**) using half of the haplotypes in the given population (AFR, EAS or EUR). These randomly drawn subsamples (each with half of the individuals in the original population) form bootstrapped samples. We define the LD of a gene pair as the average value of LD over all rare SNV pairs within that gene pair. In each iteration, we calculate the average cLD over the gene pairs in each bin (**Supplementary Materials 2.4**).

### Calculation of cLD and LD for gene pairs in 3D interaction regions

To revisit a previously negatively reported relationship between 3D interaction regions and genetic linkage disequilibrium (Whalen and Pollard 2019), we calculated both cLD and LD in a Hi-C assessment in the developing brain (Li et al.), which has 27,982 brain-specific paired 3D-interacting regions, measured from neurons derived from human induced pluripotent stem cells (hiPSCs).

Again, the 1000 Genomes Project data were used. We first calculated the distance between the genes in each pair and separate the gene pairs into 13 distance groups (**Supplementary Materials 3.1**). After stratifying all gene pairs into distance groups, within each distance group, we calculated cLD between all gene pairs and further split them into two categories: the ones that are located in 3D interaction regions (assessed by Hi-C experiments) and the ones that are located in non-3D interaction regions. The gene pairs with exactly one gene in an interaction region were discarded. Finally, the average cLD values over gene pairs within interaction and non-interaction regions were used to conduct the comparison, quantified by two two-sample tests, namely Mantel-Haenszel and Fisher exact tests (**Supplementary Materials 3.4**).

The procedure of calculating standard LD mirrors the one used above for cLD using the same distance groups and 3D-interaction vs non-interaction categories. As standard LD is defined by individual variants (not by genes), the following averaging steps were taken. For each gene pair in the 3D interaction regions, we randomly chose 2,000 rare variant pairs from it to calculate their LD values. For each selected rare variant pair, we calculated its distance and then, among the gene pairs without 3D interactions, we randomly selected another rare variant pair with the same or closest possible distance (**Supplementary Materials 3.2**). As a result, we achieved 2,000 randomly selected variant pairs from gene pairs without interaction that were matched up with the 2,000 variant pairs from gene pairs with interaction. The average values of the 2,000 variant-pairs were deemed as the LD between the gene pair.

### Calculation of cLD and LD for gene pairs in gene-gene interaction databases

Four frequently used interaction databases, Biogrid (Stark et al. 2006), Reactome (Fabregat et al. 2018), MINT (Orchard 2012) and Intact (Orchard et al. 2014) were aggregated as the source of gene-gene interactions (**Supplementary Materials 3.3**). The related datasets were downloaded from their corresponding websites and the IDs were matched using standard gene models (gencode v17). To quantify the distance between genes, only data for the gene pairs within the same chromosomes were used. Calculation of cLD and LD follows the same procedure as described for the 3D-interaction analysis, and the two-sample tests (Mantel-Haenszel and Fisher exact tests) were used to quantify the significant levels (**Supplementary Materials 3.4**).

### Protein docking analysis

We used protein docking to validate the novel gene-gene interactions predicted by unexpected high cLD values. HDOCKlite-v1.1 (Yan et al. 2020, 2017) was employed for conducting the protein-protein docking analysis between the cLD prioritized protein pairs (**Supplementary Materials 4**). The protein’s crystal structure was obtained from the Protein Data Bank (Berman et al. 2000) and further validated (Perera et al. 2021) (**Supplementary Materials 4.1**). The output file of the docked complex was visualized by PyMOL 2.5.1 (Delano), and the 2D plot of the protein-protein binding region was analyzed and deduced using LigPlot+ v.2.2 (Laskowski and Swindells 2011) (**Supplementary Materials 4.2**).

### ΔcLD genes, their functional annotation, and pathway enrichment

#### Calculation of cLD-differential gene pairs

To explore the use of cLD in distinguishing cases and controls in a typical association study, we calculated cLD using the whole exome sequencing data to study Autism Spectrum Disorder (ASD) (Satterstrom et al. 2020) [dbGaP ID: phs000298.v4.p3]. We first calculated cLD values for each gene pair for cases and controls groups separately. Then, we calculated the absolute differences between the cLD values in case and control groups for each gene pair, which was called ΔcLD. These absolute differences were sorted from largest to smallest. The top ranked genes pairs were collected and called cLD-differential gene pairs, or ΔcLD genes (**Supplementary Materials 5.2 & 5.3**).

#### Functional annotation and pathway enrichment

Based on their ΔcLD values, we selected the top 200, 500, 1,000, 1,500 and 2,000 cLD-differential gene pairs (i.e., ΔcLD genes) and used the genes sets for the downstream functional annotations. We utilized two different databases, Simons Foundation Autism Research Initiative (SFARI) (Abrahams et al. 2013) and DisGeNet (Piñero et al. 2017) as the gold-standard because they are frequently used in the field of ASD studies and general disease gene queries, respectively. We used the hypergeometric distribution probability to assess the p-value of the significance of enrichment of the cLD-differential genes against the background of gold-standard genes (**Supplementary Materials 5.4**). Additionally, using the top 2,000 cLD-differential gene pairs, we conducted GO enrichment (Ashburner et al. 2000) and KEGG pathway analysis (Kanehisa et al. 2009).

## Supporting information

cLD Supplementary Material

## Author Contributions

Conceived and supervised the study: QZ. Analyzed real data: DW, JH, DP, PK, QL. Conducted mathematical derivation and statistical simulations: DW, WZ, JW. Provided comments: CC, XG, AP. Wrote the paper: DW and QZ with major input from JH, DP, AP, and minor input from all authors.

## Data and Code Availability

The codes calculating cLD and conducting all the analyses in this work are publicly available at our GitHub: https://github.com/QingrunZhangLab/cLD

The 1000 Genome Variant Call Data used in this study could be downloaded from http://ftp.1000genomes.ebi.ac.uk. The complete variant call dataset was found using the webpage (Announcements | 1000 Genomes (internationalgenome.org)) (This is a subpage maintained by the 1000 Genome webpage) and downloaded from (Index of /vol1/ftp/release/20130502/ (ebi.ac.uk)).

The 3D Hi-C dataset is available in the Synapse database (https://www.synapse.org/) with Synapse ID: syn12979149.

The Protein Data Bank: https://www.rcsb.org/.

The DisGeNet Database: https://www.disgenet.org/

The SFARI Database: https://www.sfari.org/resource/sfari-gene/

The HDOCK protein docking software: http://hdock.phys.hust.edu.cn/

## Acknowledgments

Q.Z. is supported by NSERC Discovery Grant (RGPIN-2018-05147), University of Calgary VPR Catalyst grant and New Frontiers in Research Fund (NFRFE-2018-00748); J.W. is supported by NSERC Discovery Grant (RGPIN-2018-04328); A.P. is supported by NIH (R35 GM134957-01) and American Diabetes Association (Pathway to Stop Diabetes grant 1-19-VSN-02); D.W is supported by Alberta Graduate Excellence Scholarship; D.P. is supported by Alberta Innovates Graduate Scholarship and Eyes High International Scholarship; J.H. is supported by CSC Scholarship. The computational infrastructure is funded by Canada Foundation for Innovation JELF grant (36605).

