## Supplementary material for "cLD: Rare-variant disequilibrium between genomic regions identifies novel genomic interactions": cLD Supplementary Material

---

**SUPPLEMENTARY MATERIALS**

---

Dinghao Wang, Jingni He, Deshan Perera, Chen Cao, Pathum Kossinna,

Qing Li, William Zhang, Xingyi Guo, Alexander Platt, Jingjing Wu, Qingrun Zhang

Oct 24th 2022

### Chapter 1

### Definitions and data source

In this chapter, we mathematically defined the new statistic cLD, essentially an extension of LD analogue to the extension from MAF to cMAF. We presented its detailed mathematical formulation as well as an example. To facilitate the proofs in the next Chapter regarding the asymptotic properties of cLD (in contrast to LD), we used a mathematical language that describes the alleles and haplotypes as random variables. For this reason, LD will also be formulated using similar language and notations. We also presented details of the 1000 Genomes Project data based on which we calculated cLD and LD in various populations.

#### 1.1 Definition of LD and cLD

For a specific genomic location, we used 1 and 0 to denote the presence and absence of a non-reference allele, respectively. Then, the occurrence of a variant in a haplotype naturally follows a Bernoulli distribution, in which the success probability could be estimated by the Minor Allele Frequency (MAF) at this locus. In the example given in Supplementary Table S1.1, there are 5 individual single-SNV haplotypes $X_1, \ldots, X_5$ and each of them takes value 0 or 1.

Without losing generality, consistent to the practise in most literature, we assumed that $X_1, \ldots, X_5$ are independent and identically distributed (i.i.d.) samples drawn from

**Supplementary Table S1.1:** Example of MAF for single SNV

| POS | Gene | Info | HG1-1 | HG1-2 | HG2-1 | HG2-2 | HG3-1 | MAF |
|---|---|---|---|---|---|---|---|---|
| SNV1 | 1 | ... | $X_1$ | $X_2$ | $X_3$ | $X_4$ | $X_5$ | $p$ |

a Bernoulli distribution with success rate that is approximated by the MAF of this mutated allele.

Next we defined the scenario of cumulative case, i.e., at the region level (such as a gene), based on the genotype displayed in Supplementary Table S1.2. In this table, the shaded rows represent the genomic region for Gene 1 and the un-shaded rows are for Gene 2. $X_{ijk}$ denote whether a minor allele is present (1) or absent (0) in the $i$-th haplotype of the $j$-th gene and $k$-th variant in this gene, $i = 1, \ldots, n$ (number of haplotypes, which is twice of the number of individuals), $j = 1, \ldots, g$ (number of genes), $k = 1, \ldots, v$ (number of variants in this gene). Since $X_{ijk}$'s only take value 0 or 1, we again assume they follow Bernoulli distribution, i.e. $X_{ijk} \sim B(1, p_{jk})$, where $p_{jk}$ is the MAF of the and $k$-th variant in the $j$-th gene. $B(n, p)$ denotes the Binomial distribution with $n$ trials and $p$ being the chance of success for each trial. Note that we only considered rare variants with MAF$\leq 0.005$. The true MAF value $p_{jk}$ could be estimated by the sample MAF $\hat{p}_{jk}$. Also, we assumed that $X_{ijk}$, $i = 1, \ldots, n$, are independent random variables.

**Supplementary Table S1.2:** Cumulative effects of SNVs in a region

| POS | Gene | Info | HG1-1 | HG1-2 | HG2-1 | HG2-2 | HG3-1 | other haplotypes | MAF |
|---|---|---|---|---|---|---|---|---|---|
| 6034 | 1 | ... | $X_{111}$ | $X_{211}$ | $X_{311}$ | $X_{411}$ | $X_{511}$ | ... | $p_{11}$ |
| 6041 | 1 | ... | $X_{112}$ | $X_{212}$ | $X_{312}$ | $X_{412}$ | $X_{512}$ | ... | $p_{12}$ |
| 6047 | 1 | ... | $X_{113}$ | $X_{213}$ | $X_{313}$ | $X_{413}$ | $X_{513}$ | ... | $p_{13}$ |
| 6057 | 1 | ... | $X_{114}$ | $X_{214}$ | $X_{314}$ | $X_{414}$ | $X_{514}$ | ... | $p_{14}$ |
| 6064 | 1 | ... | $X_{115}$ | $X_{215}$ | $X_{315}$ | $X_{415}$ | $X_{515}$ | ... | $p_{15}$ |
| ... | ... | ... | ... | ... | ... | ... | ... | ... | ... |
| 7064 | 2 | ... | $X_{121}$ | $X_{221}$ | $X_{321}$ | $X_{421}$ | $X_{521}$ | ... | $p_{21}$ |
| 7124 | 2 | ... | $X_{122}$ | $X_{222}$ | $X_{322}$ | $X_{422}$ | $X_{522}$ | ... | $p_{22}$ |
| 7235 | 2 | ... | $X_{123}$ | $X_{223}$ | $X_{323}$ | $X_{423}$ | $X_{523}$ | ... | $p_{23}$ |
| ... | ... | ... | ... | ... | ... | ... | ... | ... | ... |

In order to define the cLD, with the data of rare variants in Supplementary Table S1.2, we first formed the cumulative "allele"s for each gene, illustrated in Supplementary Table S1.3. In this table, $X_{i1} = I(\sum_k X_{i1k} > 0)$ and $X_{i2} = I(\sum_k X_{i2k} > 0)$, where $I(.)$ is the logical function that will yield to 1 if the logical expression is true, and 0 otherwise. Intuitively, any SNVs being 1 will lead to the cumulative allele to be 1. The cumulative allele will be 0 only if all SNVs are 0. By counting the number of 1 in each row and dividing the total number of columns (i.e., haplotypes), one can straightforwardly calculate the cumulative MAF, or cMAF. Formally:

$$cMAF = \frac{\sum_i X_{i1}}{n}. \tag{1.1}$$

**Supplementary Table S1.3:** Cumulative genetic "allele"s for cLD calculation

| POS | Gene | Info | HG1-1 | HG1-2 | HG2-1 | HG2-2 | HG3-1 | other haplotypes |
|---|---|---|---|---|---|---|---|---|
| Gene 1 | 1 | ... | $X_{11}$ | $X_{21}$ | $X_{31}$ | $X_{41}$ | $X_{51}$ | ... |
| Gene 2 | 2 | ... | $X_{12}$ | $X_{22}$ | $X_{32}$ | $X_{42}$ | $X_{52}$ | ... |

Analogue to the traditional definition of the $r^2$ form of LD between to loci being

$$r^2 = \frac{(p_{AB} - p_A p_B)^2}{p_A(1-p_A)p_B(1-p_B)}. \tag{1.2}$$

where $A$ and $B$ are two loci, $p_{(.)}$ denotes the MAF, and the cumulative version of $p_A$, $p_B$ and $p_{AB}$ being

$$p_A = \frac{\sum_i X_{i1}}{n}, \tag{1.3}$$

$$p_B = \frac{\sum_i X_{i2}}{n}, \tag{1.4}$$

$$p_{AB} = \frac{1}{n}\sum_i X_{i1}X_{i2}. \tag{1.5}$$

we defined the cLD between Gene 1 and Gene 2 as:

$$cLD = \frac{\left[\frac{1}{n}\sum_i X_{i1}X_{i2} - \frac{1}{n^2}(\sum_i X_{i1})(\sum_i X_{i2})\right]^2}{\frac{\sum_i X_{i1}}{n}(1-\frac{\sum_i X_{i1}}{n})\frac{\sum_i X_{i2}}{n}(1-\frac{\sum_i X_{i2}}{n})}. \quad (1.6)$$

The above definition estimates the correlation between two genetic regions (e.g., genes) in terms of the aggregated effects of rare variants. In contrast, LD statistic can only be calculated for two variants. Specifically, using our notations, the LD between the $u$-th variant of Gene 1 and the $v$-th variant of Gene 2 is defined as

$$LD_{(u,v)} = \frac{\left[\frac{1}{n}\sum_i X_{i1u}X_{i2v} - \frac{1}{n^2}(\sum_i X_{i1u})(\sum_i X_{i2v})\right]^2}{\frac{\sum_i X_{i1u}}{n}(1-\frac{\sum_i X_{i1u}}{n})\frac{\sum_i X_{i2v}}{n}(1-\frac{\sum_i X_{i2v}}{n})}. \quad (1.7)$$

The above definitions of cLD and LD are based on the $r^2$ format. Alternative, one may define the cLD analogue of $D'$ below:

$$D' = \frac{D}{D_{max}} = \frac{\frac{1}{n}\sum_i X_{i1}X_{i2} - \frac{1}{n^2}(\sum_i X_{i1})(\sum_i X_{i2})}{D_{max}}$$

where

$$D = P_{AB} - P_A P_B = \frac{1}{n}\sum_i X_{i1}X_{i2} - \frac{1}{n^2}(\sum_i X_{i1})(\sum_i X_{i2}).$$

and

$$D_{\max} = \begin{cases} \max\{-P_A P_B, -(1-P_A)(1-P_B)\} \\ = \max\left\{-\frac{1}{n^2}\sum_i X_{i1}\sum_i X_{i2}, -(1-\frac{1}{n}\sum_i X_{i1})(1-\frac{1}{n}\sum_i X_{i2})\right\}, & D \leq 0, \\ \max\{P_A(1-P_B), (1-P_A)P_B\} \\ = \max\left\{\frac{1}{n}\sum_i X_{i1}(1-\frac{1}{n}\sum_i X_{i2}), (1-\frac{1}{n}\sum_i X_{i1})\frac{1}{n}\sum_i X_{i2}\right\}, & D > 0. \end{cases}$$

But in this work, we ONLY focused on the $r^2$ version, leaving the analysis of $D'$ version to the future work.

Up to now, we didn't have any assumption on the independence of variants. With these definitions, I gave the rigorous derivations of the distributions of cLD and LD in the following Chapter.

#### 1.2 An example

In this section, we used a toy example to show the calculation of cLD. The example is a small SNV dataset given in Supplementary Table S1.4, an instance of Supplementary Table S1.2.

**Supplementary Table S1.4:** Example of SNVs data for cLD calculation

| POS | Gene | Info | HG1-1 | HG1-2 | HG2-1 | HG2-2 | HG3-1 |
|---|---|---|---|---|---|---|---|
| 6034 | 1 | ... | 1 | 0 | 0 | 0 | 1 |
| 6041 | 1 | ... | 0 | 0 | 0 | 0 | 0 |
| 6047 | 1 | ... | 0 | 0 | 0 | 0 | 0 |
| 6057 | 1 | ... | 0 | 0 | 0 | 1 | 0 |
| 6064 | 1 | ... | 0 | 0 | 0 | 0 | 0 |
| 7064 | 2 | ... | 0 | 0 | 1 | 1 | 0 |
| 7124 | 2 | ... | 1 | 0 | 0 | 0 | 0 |
| 7235 | 2 | ... | 0 | 0 | 0 | 0 | 1 |

From Supplementary Table S1.4, one can form two cumulative gene alleles given in Supplementary Table S1.5 (which is an instance of the Supplementary Table S1.3 presented previously.

**Supplementary Table S1.5:** Example of cumulated gene alleles for cLD calculation

| POS | Gene | Info | HG1-1 | HG1-2 | HG2-1 | HG2-2 | HG3-1 |
|---|---|---|---|---|---|---|---|
| Gene 1 | 1 | ... | 1 | 0 | 0 | 1 | 1 |
| Gene 2 | 2 | ... | 1 | 0 | 1 | 1 | 1 |

As such, based on the formula 1.6 the cLD between Gene 1 and Gene 2 is:

$$cLD = \frac{\left[\frac{1}{n}\sum_i X_{i1}X_{i2} - \frac{1}{n^2}(\sum_i X_{i1})(\sum_i X_{i2})\right]^2}{\frac{\sum_i X_{i1}}{n}(1-\frac{\sum_i X_{i1}}{n})\frac{\sum_i X_{i2}}{n}(1-\frac{\sum_i X_{i2}}{n})} = \frac{(\frac{1}{5}\cdot 3 - \frac{1}{5^2}\cdot 3\cdot 4)^2}{0.6\cdot 0.4\cdot 0.2\cdot 0.8} = 0.375.$$

#### 1.3 Dataset and the population labels

The calculation of cLD in this paper is based on the dataset from the 1000 Genomes Project [1]. This project started in January 2008 and aimed at creating a complete and detailed catalogue of human genetic variations, which can be used for population genetic studies.

The variant call data of the Phase 3 analysis of the 1000 Genome dataset was obtained through The European Bioinformatics Institute's dedicated FTP (http://ftp.1000genomes.ebi.ac.uk) server. The complete variant call dataset was found using the webpage (Announcements — 1000 Genomes (internationalgenome.org)) (This is a sub-page maintained by the 1000 Genome webpage) and downloaded from (Index of /vol1/ftp/release/20130502/ (ebi.ac.uk)).

This dataset has 26 populations: ASW (African Ancestry in SW USA), ACB (African Caribbean in Barbados), BEB (Bengali in Bangladesh), GBR (British from England and Scotland), CDX (Chinese Dai in Xishuangbanna), CLM (Colombian in Medellín, Colombia), ESN (Esan in Nigeria), FIN (Finnish in Finland), GWD (Gambian in Western Division - Mandinka), GIH (Gujarati Indians in Houston, Texas, USA), CHB (Han Chinese in Beijing, China), CHS (Han Chinese South, China), IBS (Iberian populations in Spain), ITU (Indian Telugu in the U.K.), JPT (Japanese in Tokyo, Japan), KHV (Kinh in Ho Chi Minh City, Vietnam), LWK (Luhya in Webuye, Kenya), MSL (Mende in Sierra Leone), MXL (Mexican Ancestry in Los Angeles CA USA), PEL (Peruvian in Lima, Peru), PUR (Puerto Rican in Puerto Rico), PJL (Punjabi in Lahore, Pakistan), STU (Sri Lankan Tamil in the UK), TSI (Toscani in Italia), YRI (Yoruba in Ibadan, Nigeria), CEU (Utah Residents with Northern and Western European ancestry).

Moreover, these 26 populations could be split into five super-populations: EUR (European, 503 samples), EAS (East Asian, 504 samples), AMR (American, 347 samples), SAS (South Asian, 489 samples) and AFR (African, 661 samples). In our studies in the next chapter, we used EUR, EAS and AFR as examples to demonstrate the implementation and performance of cLD. The selection was based on that they are frequently used in other literature and they do not contain substantial population admixtures. Relatives have been removed to ensure that samples are generally unrelated. More specifically, if an individual's parent or sibling is present in the dataset, then this person is removed from the calculation.

### Chapter 2

### Asymptotic Properties of cLD

In this chapter, we derived the asymptotic distributions of cLD and LD to show that cLD enjoys lower variance and is more stable than LD. This has been done using two distinct mathematical approaches, outlined below:

First, we used colsed-form derivation of the variance of LD and cLD and used them to calculate their variances using parameters collected from the 1000 Genomes Project data. More specifically, we first modelled the exact distribution of cLD and LD via multinomial random variables using the allele counts, and then used multivariate Central Limit Theory (CLT) and multivariate Delta method to derive the asymptotic distributions of LD and cLD respectively. These results are presented in Sections 2.1 and 2.2. Such results allow us to derive the variances in closed form ready to be calculated. In Section 2.3, we compared the variability of cLD with LD and showed that cLD has a lower variance than LD.

As the above derivation assumes large samples (for the multivariate CLT to be valid), for the cases of limited samples, in section 2.4, we used bootstrap method to show that cLD is more stable than LD in the partial samples. Here the property of being "sable" is reflected by the slimmer distribution (i.e., with lower variability) formed by the bootstrapped samples.

#### 2.1 Derivation of closed-form distribution of LD

##### 2.1.1 Multinomial distribution

With the same notation defined in Chapter 1, we used $X_{ijk}$ to denote the $i$-th haplotype of the $j$-th gene and the $k$-th variant. Without loss of generality, we assumed that SNV1 and SNV2 come from Gene 1 and Gene 2 respectively. The data for two SNVs are denoted in Supplementary Table S2.1. In such a case, each haplotype has a pair of SNVs (SNV1 is the $u$-th variant from Gene 1 while SNV2 is the $v$-th variant from Gene 2), and the $i$-th pair $(X_{i1u}, X_{i2v})$, $i = 1, \ldots, n$, on (SNV1, SNV2) can take possible values $(1, 0)$, $(1, 1)$, $(0, 1)$ and $(0, 0)$. Further, these $n$ pairs can be treated as i.i.d. following a multinomial distribution given in Supplementary Table S2.2. This table shows a multinomial distribution for the $i$-th haplotype, where $O_i$'s denote the frequency for each possible pair value. The third column lists the counts for each possible outcome of the pair $(X_{i1u}, X_{i2v})$ throughout the entire dataset, and the fourth column represents the corresponding population probabilities of the multinomial distribution. Intuitively the population parameters $p_1, \ldots, p_4$ can be estimated by sample fractions, i.e.

$$\hat{p}_i = O_i/n, \quad i = 1, \ldots, 4.$$

**Supplementary Table S2.1:** Data of two SNVs

| POS | Gene | Info | HG01 | HG02 | HG03 | HG04 | HG05 | ... | MAF |
|---|---|---|---|---|---|---|---|---|---|
| SNV1 | 1 | ... | $X_{11u}$ | $X_{21u}$ | $X_{31u}$ | $X_{41u}$ | $X_{51u}$ | ... | $p_{11}$ |
| SNV2 | 2 | ... | $X_{12v}$ | $X_{22v}$ | $X_{32v}$ | $X_{42v}$ | $X_{52v}$ | ... | $p_{21}$ |

**Supplementary Table S2.2:** Multinomial modelling of two SNVs

| $X_{i1u}$ | $X_{i2v}$ | Count | Probability |
|---|---|---|---|
| 1 | 1 | $O_1$ | $p_1$ |
| 0 | 1 | $O_2$ | $p_2$ |
| 1 | 0 | $O_3$ | $p_3$ |
| 0 | 0 | $O_4 = n - O_1 - O_2 - O_3$ | $p_4 = 1 - p_1 - p_2 - p_3$ |

With these preparations of defining notations, we estimated the LD $((r^2))$ and then derive the asymptotic distribution of the resulted estimator. Firstly, the LD can be rewritten in terms of current parameters as

$$LD_{(u,v)} = \left[\frac{p_{AB} - p_A p_B}{\sqrt{p_A(1-p_A)p_B(1-p_B)}}\right]^2 = \left[\frac{p_1 - (p_1+p_3)(p_1+p_2)}{\sqrt{(p_1+p_3)(p_2+p_4)(p_1+p_2)(p_3+p_4)}}\right]^2. \quad (2.1)$$

Then the LD can be estimated by plugging-in the estimators using the counts $O_x$, $(x = 1, 2, 3, 4)$:

$$\begin{aligned} \widehat{LD}_{(u,v)} &= \left[\frac{\hat{p}_1 - (\hat{p}_1+\hat{p}_3)(\hat{p}_1+\hat{p}_2)}{\sqrt{(\hat{p}_1+\hat{p}_3)(\hat{p}_2+\hat{p}_4)(\hat{p}_1+\hat{p}_2)(\hat{p}_3+\hat{p}_4)}}\right]^2 \\ &= \frac{\left[\frac{O_1}{n} - \frac{1}{n^2}(O_1+O_2)(O_1+O_3)\right]^2}{\frac{O_1+O_3}{n}(1-\frac{O_1+O_3}{n})\frac{O_1+O_2}{n}(1-\frac{O_1+O_2}{n})} \\ &= \frac{(O_1O_4 - O_2O_3)^2}{(O_1+O_2)(O_1+O_3)(O_2+O_4)(O_3+O_4)}. \end{aligned} \quad (2.2)$$

Though both LD and $\widehat{LD}$ are calculated for fixed variant pair $(u, v)$, for notation simplicity we remove their dependence on $(u, v)$ in the subscripts and let $LD(\boldsymbol{p}) = LD_{(u,v)}$ and $LD(\boldsymbol{O}/n) = \widehat{LD}_{(u,v)}$, where $\boldsymbol{p} = (p_1, p_2, p_3, p_4)^\top$ and $\boldsymbol{O} = (O_1, O_2, O_3, O_4)^\top$.

The mean of the estimator $LD(\boldsymbol{O}/n)$ is thus given by:

$$E[LD(\boldsymbol{O}/n)] = \sum_{\Omega_n} \frac{(O_1O_4 - O_2O_3)^2}{(O_1+O_2)(O_1+O_3)(O_2+O_4)(O_3+O_4)} \cdot p(O_1, O_2, O_3),$$

where $\Omega_n = \{(O_1, O_2, O_3) : O_i \in \mathbb{N} \cup \{0\}, O_i \leq n, O_1 + O_2 + O_3 \leq n\}$ and

$$p(O_1, O_2, O_3) = \frac{n!}{O_1!O_2!O_3!O_4!} {p_1}^{O_1} {p_2}^{O_2} {p_3}^{O_3} {p_4}^{O_4}.$$

The variance of $LD(\boldsymbol{O}/n)$ is $Var(LD(\boldsymbol{O}/n)) = E\left[(LD(\boldsymbol{O}/n))^2\right] - E^2[LD(\boldsymbol{O}/n)]$.

It is known that when sample size $n$ is reasonably large, the calculation of multinomial probabilities are very tedious, thus the expressions for the mean and variance of $LD(\boldsymbol{O}/n)$

given above are not practical. To solve this problem, we next used the multivariate CLT and Delta method to approximate the exact distribution of $LD(\boldsymbol{O}/n)$ when $n$ is large.

##### 2.1.2 Asymptotic normality approximation of LD

For notation simplicity, we defined events $E_1$ as $(X_{1u}, X_{2v}) = (1,1)$ on (SNV1, SNV2), $E_2$ as $(X_{1u}, X_{2v}) = (0,1)$ on (SNV1, SNV2), $E_3$ as $(X_{1u}, X_{2v}) = (1,0)$ on (SNV1, SNV2), and $E_4$ as $(X_{1u}, X_{2v}) = (0,0)$ on (SNV1, SNV2). Then

$$\boldsymbol{O} = \begin{pmatrix} O_1 \\ O_2 \\ O_3 \\ O_4 \end{pmatrix} = \sum_{i=1}^{n} \begin{pmatrix} I(E_1 \text{ occurs on the } i\text{−th haplotype}) \\ I(E_2 \text{ occurs on the } i\text{−th haplotype}) \\ I(E_3 \text{ occurs on the } i\text{−th haplotype}) \\ I(E_4 \text{ occurs on the } i\text{−th haplotype}) \end{pmatrix} = \sum_{i=1}^{n} \boldsymbol{I_i},$$

where $\boldsymbol{I_i} = (I_{i1}, I_{i2}, I_{i3}, I_{i4})^\top$ and

$$I_{ij} = I(E_j \text{ occurs on the } i\text{−th haplotype}), \quad j = 1,2,3,4,\ i = 1,\ldots,n.$$

As a result, $\boldsymbol{O} \sim Multinom(n; \boldsymbol{p})$ with $\boldsymbol{p} = (p_1, p_2, p_3, p_4)^\top$. Since these $n$ samples are independent, by multivariate central limit theorem (CLT), we have the asymptotic normality of $\frac{\boldsymbol{O}}{n}$ given by

$$\sqrt{n}\left(\frac{\boldsymbol{O}}{n} - \boldsymbol{p}\right) \xrightarrow{\mathcal{L}} N(\boldsymbol{0}, \Sigma),$$

where $\xrightarrow{\mathcal{L}}$ denotes convergence in distribution and the asymptotic covariance matrix $\Sigma = Cov(\boldsymbol{I_1}) = (\sigma_{jl})$. Straightforward calculation gives

$$\sigma_{jl} = E[I_{1j}I_{1l}] - E[I_{1j}]E[I_{1l}] = P(I_{1j} = I_{1l} = 1) - p_j p_l = \begin{cases} -p_j p_l, & j \neq l, \\ p_j - p_j p_l, & j = l. \end{cases}$$

Thus

$$\Sigma = \begin{pmatrix} p_1 - p_1^2 & -p_1p_2 & -p_1p_3 & -p_1p_4 \\ -p_2p_1 & p_2 - p_2^2 & -p_2p_3 & -p_2p_4 \\ -p_3p_1 & -p_3p_2 & p_3 - p_3^2 & -p_3p_4 \\ -p_4p_1 & -p_4p_2 & -p_4p_3 & p_4 - p_4^2 \end{pmatrix} = \boldsymbol{D_p} - \boldsymbol{p}\boldsymbol{p}^\top, \tag{2.3}$$

where $\boldsymbol{D_p}$ is the diagonal matrix with diagonal elements $p_1, \ldots, p_4$. Note that $\Sigma$ is singular, since the sum of each row of $\Sigma$ gives 0. As a result, when sample size $n$ is large, we have the approximated distributions of $\boldsymbol{O}/n$ and $\boldsymbol{O}$ given by

$$\frac{\boldsymbol{O}}{n} \sim AN\left(\boldsymbol{p}, \frac{\Sigma}{n}\right), \quad \boldsymbol{O} \sim AN\left(n\boldsymbol{p}, n\Sigma\right),$$

where 'AN' stands for Asymptotic Normal.

In order to derive the asymptotic distributions of cLD and LD, first note that the LD given in (1.7) is a function of $\boldsymbol{O}$, thus the multivariate Delta method applied to the asymptotic normality of $\boldsymbol{O}$ would give us the asymptotic normality of LD. Let $\boldsymbol{J}_{LD} = \left[\frac{\partial LD(\boldsymbol{O}/n)}{\partial \boldsymbol{O}}\right]\Big|_{\boldsymbol{O}=n\boldsymbol{p}}$ be the $1\times 4$ Jacobian Matrix of $LD(\boldsymbol{O}/n)$, with the $(1, j)$-th element being $\left(\frac{\partial LD(\boldsymbol{O}/n)}{\partial O_j}\right)|_{\boldsymbol{O}=n\boldsymbol{p}}$, where

$$\frac{\partial LD(\boldsymbol{O}/n)}{\partial O_1} = \frac{(O_1O_4 - O_2O_3)[2O_2O_3(O_1+O_4) + (O_2+O_3)(O_1O_4+O_2O_3)]}{(O_1+O_2)^2(O_1+O_3)^2(O_2+O_4)(O_3+O_4)},$$

$$\frac{\partial LD(\boldsymbol{O}/n)}{\partial O_2} = -\frac{(O_1O_4 - O_2O_3)[2O_1O_4(O_2+O_3) + (O_1+O_4)(O_1O_4+O_2O_3)]}{(O_1+O_2)^2(O_1+O_3)(O_2+O_4)^2(O_3+O_4)},$$

$$\frac{\partial LD(\boldsymbol{O}/n)}{\partial O_3} = -\frac{(O_1O_4 - O_2O_3)[2O_1O_4(O_2+O_3) + (O_1+O_4)(O_1O_4+O_2O_3)]}{(O_1+O_2)(O_1+O_3)^2(O_2+O_4)(O_3+O_4)^2},$$

$$\frac{\partial LD(\boldsymbol{O}/n)}{\partial O_4} = \frac{(O_1O_4 - O_2O_3)[2O_2O_3(O_1+O_4) + (O_2+O_3)(O_1O_4+O_2O_3)]}{(O_1+O_2)(O_1+O_3)(O_2+O_4)^2(O_3+O_4)^2}.$$

Then multivariate Delta method gives the asymptotic distribution of $LD(\boldsymbol{O}/n)$ as

$$LD(\boldsymbol{O}/n) - LD(\boldsymbol{p}) \sim AN\left(0, n\boldsymbol{J}_{LD}\Sigma\boldsymbol{J}_{LD}^\top\right), \tag{2.4}$$

where $\Sigma$ is given in (2.3).

The derived asymptotic distribution of $LD(\boldsymbol{O}/n)$ given in (2.4) can be used for various downstream tasks. For instance, one can assess the significance of LD using the $p$-value calculated by the distribution.

In this work, the use of the above distribution is to estimate the asymptotic variance of $LD(\boldsymbol{O}/n)$, $n\boldsymbol{J}_{LD}\Sigma\boldsymbol{J}_{LD}^{\top}$, which can not be used directly in practice due to the unknown $\boldsymbol{p}$ involved. Thus to estimate $Var(\widehat{LD})$, intuitively we have to use plug-in method by replacing the unknowns by the observed allele counts. More specifically, we let $\widehat{\Sigma}$ denote the $\Sigma$ in (2.3) with $\boldsymbol{p}$ replaced by $\boldsymbol{O}/n$ and let $\widehat{\boldsymbol{J}}_{LD} = \frac{\partial LD(\boldsymbol{O}/n)}{\partial \boldsymbol{O}}$, then when sample size $n$ is large, the variance of $LD(\boldsymbol{O}/n)$ can be estimated by

$$\widehat{Var}(LD(\boldsymbol{O}/n)) = n\widehat{\boldsymbol{J}}_{LD}\widehat{\Sigma}\widehat{\boldsymbol{J}}_{LD}^{\top}. \tag{2.5}$$

#### 2.2 Derivation of closed-form distribution of cLD

The asymptotic distribution and estimated variance of cLD could be similarly derived as to those of LD in Section 2.1. The difference between cLD and LD is that they are functions of different data. LD is calculated on two SNVs, while cLD is calculated on aggregated effect of SNVs in two genes. That said, the forms of SNV pairs and gene pairs are mathematically similar and can be modelled similarly. Here, we extended the derivations for LD to cLD with some non-essential modifications.

For $n$ independent pairs $(X_{i1}, X_{i2})$, $i = 1, \ldots, n$, observed on two genes (Gene 1, Gene 2), they can be treated as i.i.d. following a multinomial distribution given in Supplementary Table S2.3, along with frequencies $M_i$'s for each possible pair value.

Firstly, the cLD ($r^2$) can be rewritten in terms of current parameters as

$$cLD := cLD(\boldsymbol{q}) = \left[\frac{q_1 - (q_1 + q_3)(q_1 + q_2)}{\sqrt{(q_1 + q_3)(q_2 + q_4)(q_1 + q_2)(q_3 + q_4)}}\right]^2. \tag{2.6}$$

**Supplementary Table S2.3:** Multinomial modelling of two gene lines

| $X_{i1}$ | $X_{i2}$ | Count | Probability |
|---|---|---|---|
| 1 | 1 | $M_1$ | $q_1$ |
| 0 | 1 | $M_2$ | $q_2$ |
| 1 | 0 | $M_3$ | $q_3$ |
| 0 | 0 | $M_4 = n - M_1 - M_2 - M_3$ | $q_4 = 1 - q_1 - q_2 - q_3$ |

Then the cLD can be estimated by the plug-in estimator

$$\begin{aligned} \widehat{cLD} := cLD(\boldsymbol{M}/n) &= \frac{\left[\frac{M_1}{n} - \frac{1}{n^2}(M_1+M_2)(M_1+M_3)\right]^2}{\frac{M_1+M_3}{n}(1-\frac{M_1+M_3}{n})\frac{M_1+M_2}{n}(1-\frac{M_1+M_2}{n})} \\ &= \frac{(M_1M_4 - M_2M_3)^2}{(M_1+M_2)(M_1+M_3)(M_2+M_4)(M_3+M_4)}. \end{aligned} \tag{2.7}$$

where $\boldsymbol{M} = (M_1, M_2, M_3, M_4)^\top$ and $\boldsymbol{q} = (q_1, q_2, q_3, q_4)^\top$. Following exactly the same steps as in Section 2.1 for LD, we have

$$cLD(\boldsymbol{M}/n) - cLD(\boldsymbol{q}) \sim AN\left(0, n\boldsymbol{J}_{cLD}\boldsymbol{Q}\boldsymbol{J}_{cLD}^\top\right), \tag{2.8}$$

where

$$\boldsymbol{Q} = \begin{pmatrix} q_1 - q_1^2 & -q_1q_2 & -q_1q_3 & -q_1q_4 \\ -q_2q_1 & q_2 - q_2^2 & -q_2q_3 & -q_2q_4 \\ -q_3q_1 & -q_3q_2 & q_3 - q_3^2 & -q_3q_4 \\ -q_4q_1 & -q_4q_2 & -q_4q_3 & q_4 - q_4^2 \end{pmatrix}, \tag{2.9}$$

$$\boldsymbol{J}_{cLD} = \left[\frac{\partial cLD(\boldsymbol{M}/n)}{\partial \boldsymbol{M}}\right]\Bigg|_{\boldsymbol{M}=n\boldsymbol{q}} = \left(\frac{\partial cLD(\boldsymbol{M}/n)}{\partial M_1}, \frac{\partial LD(\boldsymbol{M}/n)}{\partial M_2}, \frac{\partial cLD(\boldsymbol{M}/n)}{\partial M_3}, \frac{\partial cLD(\boldsymbol{M}/n)}{\partial M_4}\right)\Bigg|_{\boldsymbol{M}=n\boldsymbol{q}}.$$

Similar to the situation in the last section, from (2.8) we see that the asymptotic variance of $cLD(\boldsymbol{M}/n)$ is $n\boldsymbol{J}_{cLD}\boldsymbol{Q}\boldsymbol{J}_{cLD}^\top$, which can not be used directly in practice due to the unknown $\boldsymbol{q}$ involved. Thus to estimate $Var(\widehat{cLD})$, we let $\widehat{\boldsymbol{Q}}$ denote the $\boldsymbol{Q}$ in (2.9) with $\boldsymbol{q}$ replaced by $\boldsymbol{M}/n$ and let $\widehat{\boldsymbol{J}}_{cLD} = \frac{\partial cLD(\boldsymbol{M}/n)}{\partial \boldsymbol{M}}$, then when sample size $n$ is large, the variance of

$\widehat{cLD} = cLD(\boldsymbol{M}/n)$ can be estimated by

$$\widehat{Var}(cLD(\boldsymbol{M}/n)) = n\widehat{\boldsymbol{J}}_{cLD}\widehat{\boldsymbol{Q}}\widehat{\boldsymbol{J}}_{cLD}^{\top}. \tag{2.10}$$

#### 2.3 Comparison of variances using closed-form distributions

In Section 2.1 and 2.2, we have derived the asymptotic variances of $cLD(\boldsymbol{M}/n)$ and $LD(\boldsymbol{O}/n)$ and provided their variance estimates which can be calculated directly from data. In this section, we conducted the comparison between cLD and LD in terms of their variability by plugging-in the frequency estimates using the 1000 Genomes Project data.

First, we randomly sampled 1000 gene pairs from the 1000 Genomes Project data. Second, we calculated $Var(cLD(\boldsymbol{M}/n))$ for each gene pair by their estimated cMAF in the real data; and then obtain the average value $\overline{Var}(cLD(\boldsymbol{M}/n))$ over all gene pairs in our sample. Third, similarly, within each gene pair, we calculated the $Var(LD(\boldsymbol{O}/n))$ for each rare SNV pair, and we define the variance of LD of a gene pair as the average value of $Var(LD(\boldsymbol{O}/n))$ over all rare SNV pairs within that gene pair. As such, we obtained the average value of variance $\overline{Var}(cLD(\boldsymbol{M}/n))$ and $\overline{Var}(LD(\boldsymbol{O}/n))$ over the randomly sampled gene pairs.

However, as cLD is not at the same scale as LD, the direct comparison between the variances may be unfair. In the above setting, as there are many pairs of SNVs between a pair of genes (therefore many LD values) whereas only one cLD between a pair of genes, the LD has been highly averaged, leading to lower variance. In contrast, if we do not conduct such within-gene average, and only conduct the sum or taking a random pair of SNVs, it is unfair for LD.

To elaborate the above intuition mathematically, let us look at the formula of variances:

$$Var(X) = E[X - E(X)]^2$$

where X is a random variable. If the scale of X is a small value, then the variance could also be also small. Thus, in order to compare the stability of them, we mush use mean value to re-scale $cLD(\boldsymbol{M}/n)$ and $LD(\boldsymbol{O}/n)$.

Based on the above rationale, we used the following formulas to conduct the re-scaling before the comparison:

$$cLD_{scaled} = \frac{cLD(\boldsymbol{M}/n)}{mean\ value\ of\ cLD(\boldsymbol{M}/n)}$$

and

$$LD_{scaled} = \frac{LD(\boldsymbol{O}/n)}{mean\ value\ of\ LD(\boldsymbol{O}/n)}.$$

Where the mean values of LD and cLD are calculated over all gene pairs in our sample. Thus we can compare the scaled variance of LD and cLD:

$$\overline{Var}(cLD)_{scaled} = \frac{\overline{Var}(cLD(\boldsymbol{M}/n))}{(mean\ value\ of\ cLD(\boldsymbol{M}/n))^2}$$

and

$$\overline{Var}(LD)_{scaled} = \frac{\overline{Var}(LD(\boldsymbol{O}/n)))}{(mean\ value\ of\ LD(\boldsymbol{O}/n))^2}.$$

The result from 1000 Genomes Project data shows:

$$\overline{Var}(cLD)_{scaled} \approx 1.3 \times 10^8 \ll 3.6 \times 10^{14} \approx \overline{Var}(LD)_{scaled} \qquad (2.11)$$

Evidently, the variance of cLD is **six** magnitude lower than the one of LD, indicating it is a way stabler statistic than standard LD.

Moreover, we wanted to stratify the comparison of variance by the value of cMAF. To do this, we split the gene pairs into 4 groups according to the smaller cMAF of the two genes (in the pair to calculate LD and cLD). The cMAF intervals are set to be [0, 0.05], [0.05, 0.1], [0.1, 0.2] and [0.2, 0.4]. After separating gene pairs into 4 groups, we calculated the average variance of cLD using the closed-form formula over gene pairs for each group. Thus we have 4 values for Var(cLD), matching the 4 cMAF groups. Similarly, for each cMAF group, we used the same method in this section to calculate the average Var(LD). Finally, we calculate the ratio for each cMAF group:

$$Ratio \ = \ \frac{\overline{Var}(cLD)}{\overline{Var}(LD)}$$

The results of EUR, AFR and EAS population are shown in Supplementary Fig. S2.1 (a), Supplementary Fig. S2.2(a) and Supplementary Fig. S2.3(a), respectively. They show the same pattern that, in all populations, cLD is more stable at all cMAF spectrum, with the most significant effect at the low cMAF end.

#### 2.4 Estimating stability by Bootstrapped distributions

In Section 2.3, we used the closed-form formula to compare the variance of scaled cLD and LD, and concluded that cLD is stabler. To conduct such comparisons thoroughly, in this section, we used a different strategy, the Bootstrap method, to compare the variance of cLD and LD. We are still using the 1000 Genomes Project data in the calculation.

The idea of this comparison is that we randomly sampled half of the haplotypes in a given population and calculated the average cLD and average LD over the gene pairs within a cMAF group, and then we repeat this procedure 500 times. After that, we have 500 values of cLD and LD. Base on these cLDs and LDs from 500 iterations, we can use the same method in Section 2.3 to re-scale the cLD and LD, and then calculate the distribution of scaled cLD and LD. In order to study the effect of cMAF on the distribution, we separate the gene pairs into 10 groups according to the two cMAFs of the two genes in the gene pair. More specifically, the steps are as follows.

Step 1: Randomly sample 1000 genes from the 1000 Genomes Project data.

Step 2: Separate genes into 4 bins according to these threshold:

$$Bin\ 1:\ cMAF\ in\ [0.2,\ 0.4]$$
$$Bin\ 2:\ cMAF\ in\ [0.1,\ 0.2]$$
$$Bin\ 3:\ cMAF\ in\ [0.05,\ 0.1]$$
$$Bin\ 4:\ cMAF\ in\ [0,\ 0.05]$$

Then pair-wisely, the genes were paired into 10 groups:

$$(Bin\ 1, Bin\ 1),\ (Bin\ 1, Bin\ 2),\ (Bin\ 1, Bin\ 3),\ (Bin\ 1, Bin\ 4),$$
$$(Bin\ 2, Bin\ 2),\ (Bin\ 2, Bin\ 3),\ (Bin\ 2, Bin\ 4),$$
$$(Bin\ 3, Bin\ 3),\ (Bin\ 3, Bin\ 4),\ and$$
$$(Bin\ 4, Bin\ 4).$$

Then we repeated the following Step 3 to Step 5 for 500 times.

Step 3: Randomly sampled half of the haplotypes in the given population (AFR, EAS or EUR).

Step 4: For each gene pair, we calculated the cLD and LD. Similar to Section 2.3, we define the LD of a gene pair as the average value of $LD(\boldsymbol{O}/n)$ over all rare SNV pairs within that gene pair.

Step 5: In each iteration, calculated the avarage cLD (denoted as $cLD_{ij}$) and LD (denoted as $LD_{ij}$) over the gene pairs in each group, where i denotes the i-th iteration (i = 1, 2, ..., 500) and j denotes the group number (j = 11, 12 ,13, 14, 22, 23, 24, 33, 34, 44).

Step 6: Lastly, base on $cLD_{ij}$ and $LD_{ij}$ (i = 1,2...500 and j = 11, 12 ,13, ..., 44), we obtained the density of cLD and LD for 10 cMAF groups. To be more specific, we are using kernel estimation to smooth the distribution for the visualization.

The results of EUR, AFR and EAS population are shown in Supplementary Fig. S2.1 (b), Supplementary Fig. S2.2 (b) and Supplementary Fig. S2.3 (b), respectively. These results show that cLD has a way slimmer bootstrapped distribution than LD, further evidencing the stability of cLD over LD.

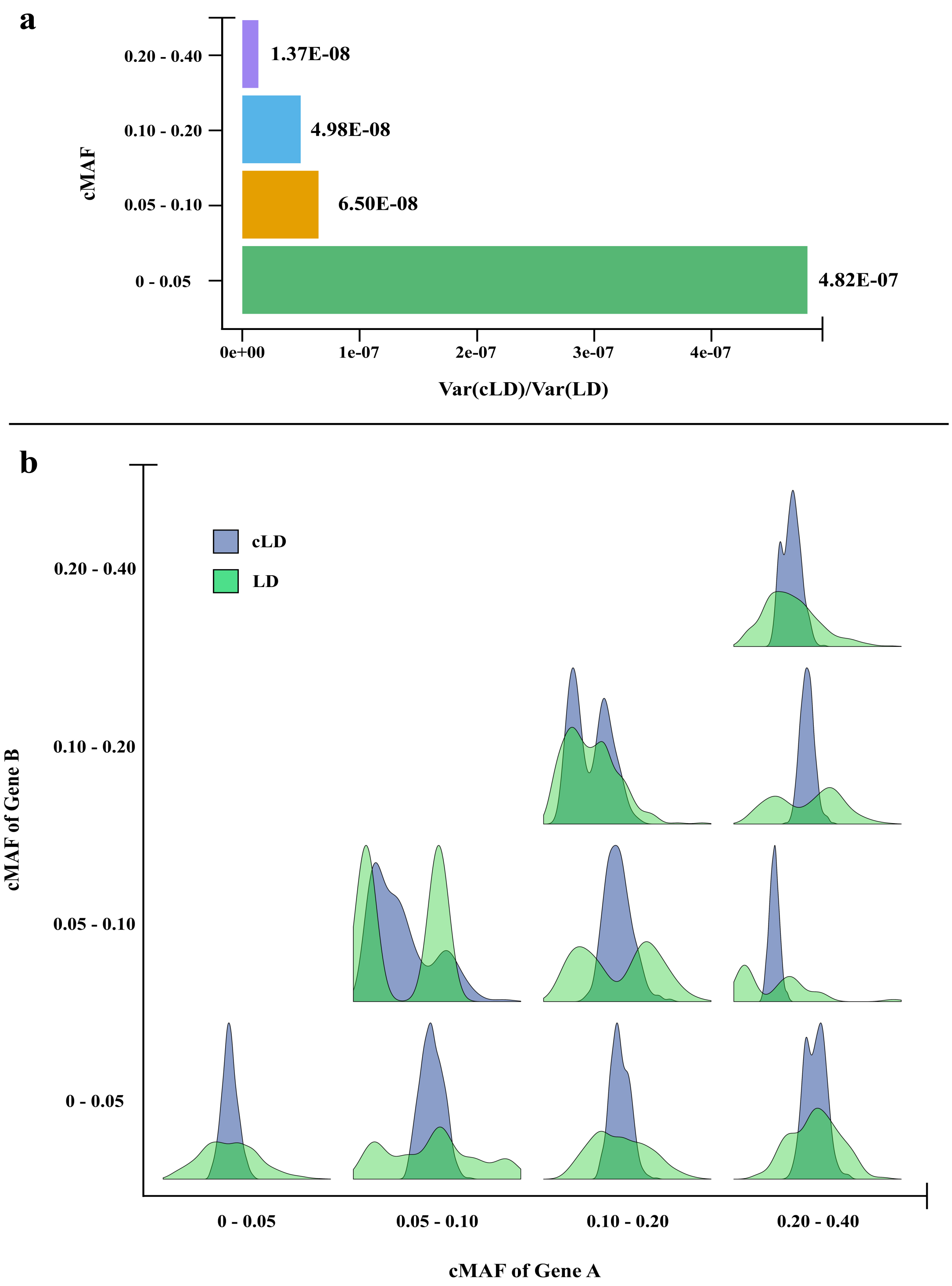

**Supplementary Fig. S2.1:** Stability of cLD and LD revealed by closed-form variance calculation and bootstrapped resampling. The EUR population is shown. a) The gene pairs were split into four different bins based on the cMAF values, i.e., $< 0.05$, 0.05 - 0.1, 0.1 - 0.2, and 0.2-0.4, which has been shown in the y-axis. The x-axis is the ratio between the variance of LD and cLD. b) Probability density distribution of cLD and LD by generating bootstrapped samples.

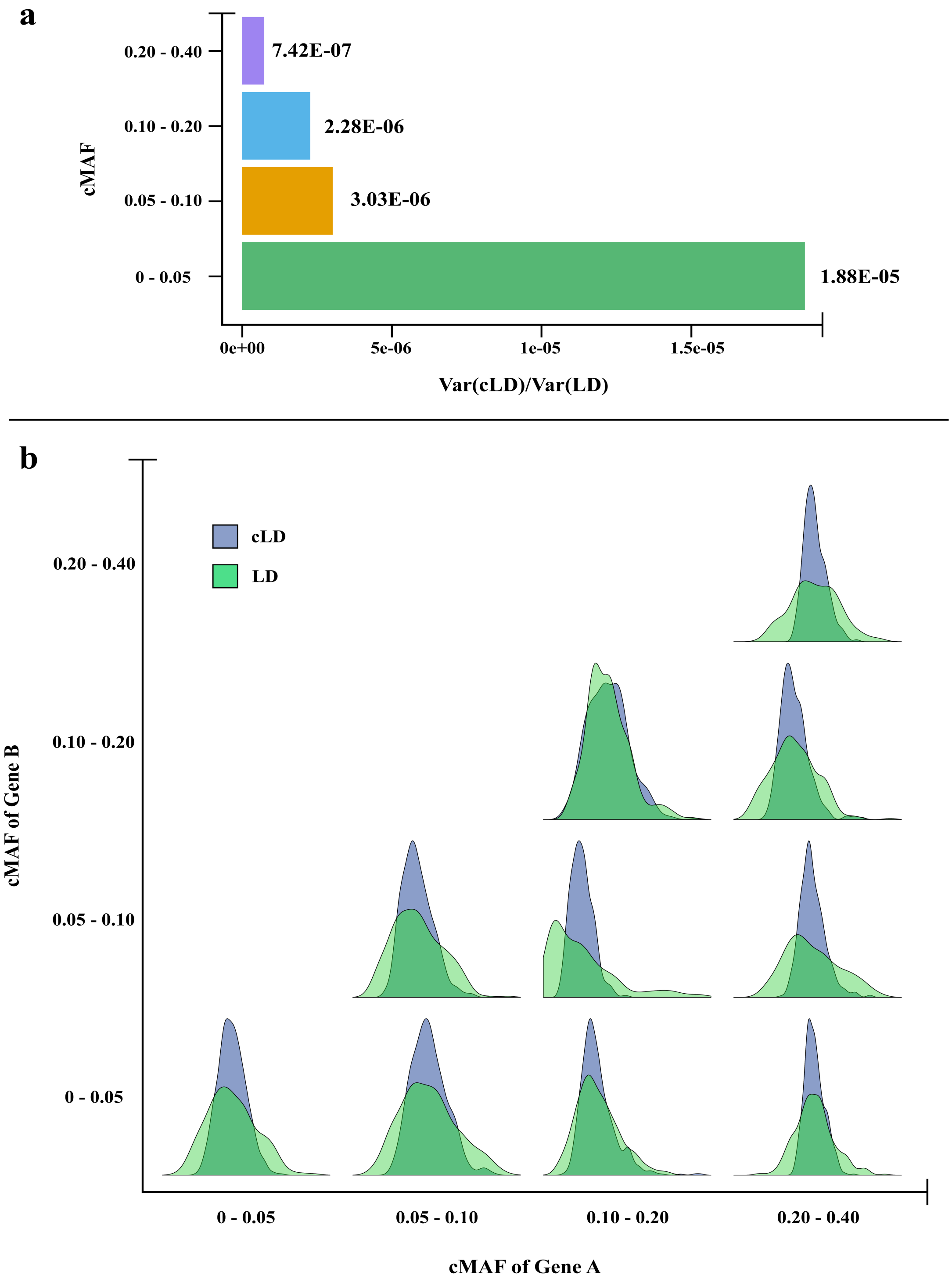

**Supplementary Fig. S2.2:** Stability of cLD and LD revealed by closed-form variance calculation and bootstrapped resampling. The AFR population is shown. a) The gene pairs were split into four different bins based on the cMAF values, i.e., $< 0.05$, 0.05 - 0.1, 0.1 - 0.2, and 0.2-0.4, which has been shown in the y-axis. The x-axis is the ratio between the variance of LD and cLD. b) Probability density distribution of cLD and LD by generating bootstrapped samples.

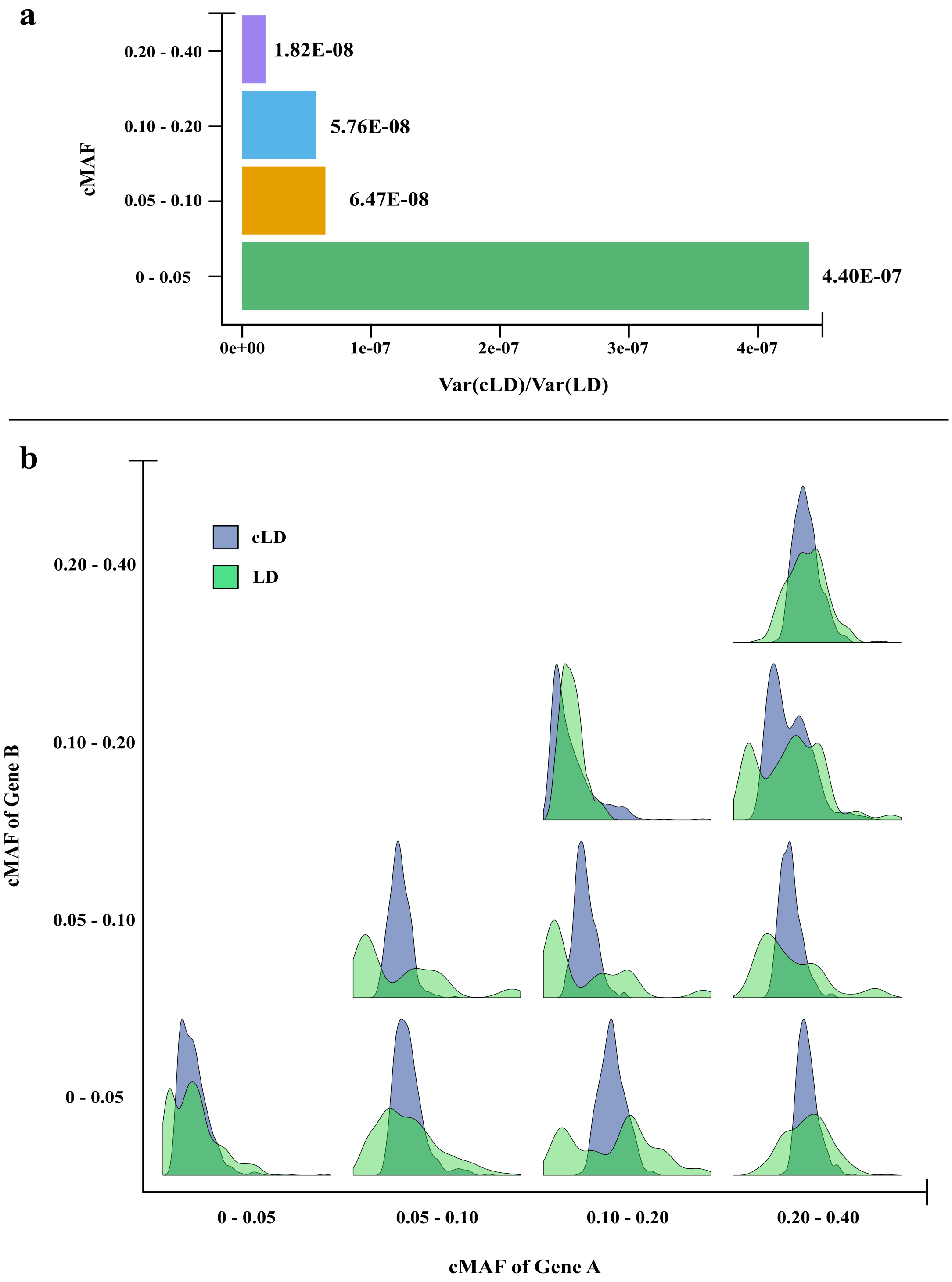

**Supplementary Fig. S2.3:** Stability of cLD and LD revealed by closed-form variance calculation and bootstrapped resampling. The EAS population is shown. a) The gene pairs were split into four different bins based on the cMAF values, i.e., < 0.05, 0.05 - 0.1, 0.1 - 0.2, and 0.2-0.4, which has been shown in the y-axis. The x-axis is the ratio between the variance of LD and cLD. b) Probability density distribution of cLD and LD by generating bootstrapped samples.

### Chapter 3

### Applying cLD to real sequence data

In this chapter, to show the practical use of cLD in capturing genetic interactions, we calculated cLD using phased version of the 1000 Genomes Project data described in Chapter 1 and characterized its enrichment based on known interactions. These interactions include both 3D spatial genomic interactions (assessed by the Hi-C experiments) and the four popular interaction databases.

The rest of this chapter is organized as follows. Because such disequilibrium statistics (including both cLD and LD) in general decrease as the distance between the two loci increases, in Section 3.1, we defined the bins according to the distance between the pairs as the units of the analyses in this chapter. We then use cLD to assess the enrichment of high cLD pairs in the 3D interacting regions. In contrast to the negative results reported previously using standard LD [2], we indeed discovered the higher cLD in 3D interaction regions in Section 3.2. Next, we applied cLD to four interaction databases Intact, Reactom, MINT & Biogrid and discovered that cLD is also enriched in gene pairs with known interactions in Section 3.3. To rigorously assess the statistical significance of the above mentioned enrichment in both 3D and gene-gene interactions, we designed the use of the Fisher's Exact test and Mantel-Haenszel test to quantify the p-values of the comparisons in the previous sections. Finally, in Section 3.5, we provided the details of enrichment analysis that shows the higher proportion of pairs of genes in 3D interaction regions.

#### 3.1 Gene pairs grouped by distance bins

To give one single number of the location of a gene, we calculated the mean value of the positions of all SNVs in this gene. Then for every pairs of genes, the difference of their locations (as calculated above) will be their distance. Based on the distances of the gene pairs, they are split in to 13 groups using the cut points below:

$35 \cdot 10^3 = 35Kb$, $70 \cdot 10^3 = 70Kb$, $140 \cdot 10^3 = 140Kb$, $280 \cdot 10^3 = 280Kb$, $560 \cdot 10^3 = 560Kb$, $1.12 \cdot 10^6 = 1.12Mb$, $2.24 \cdot 10^6 = 2.24Mb$, $4.48 \cdot 10^6 = 4.48Mb$, $8.96 \cdot 10^6 = 8.96Mb$, $17.92 \cdot 10^6 = 17.92Mb$, $35.84 \cdot 10^6 = 35.84Mb$, $71.6 \cdot 10^6 = 71.6Mb$ and $143.2 \cdot 10^3 = 143.2Mb$.

The above grouping applies to all the analysis in this chapter.

#### 3.2 LD and cLD in 3D interacting regions assessed by Hi-C experiments

There are a series of emerging molecular biology methods for analyzing the spatial organization of chromatin in a cell using next-generation sequencing instruments. Using these techniques, researchers can quantify the interactions between loci that are nearby in 3D space, but may not be close to each other in the linear genome [3]. Among these techniques, Hi-C is a frequently used techniques assessing 3D structure in a high resolution.

There was a widely spread expectation that the 3D genomic interaction in the form of chromatin contact may leave a footprint in the form of genetic LD. Motivated by such discussions, Whalen and Pollard have measured the LD using $r^2$ based on the common variants (MAF $> 0.05$) in 1000 Genomes Project data and reported negative results stating that genetic LD maps are not overlapping with the 3D contact map. However, in this section, by re-analyzing the same genotype data and the Hi-C revealed 3D contact map in the developing brain using cLD and rare variants, we revealed that the 3D chromatin interactions did leave genetic footprints in the form of higher cLD in pairs of genes that are in the contacting Hi-C regions.

Since cLD statistic is designed to capture the association between genes, we expect the cLD between gene pairs that are in the 3D-interaction regions to be higher than the cLD between gene pairs that do not. To demonstrate this, we compare the average cLD of gene pairs in 3D-interaction regions against the average cLD of gene-pairs not in 3D-interaction regions. For an appropriate comparison, we stratify the data using the different distance bins defined in section 3.1.

The distribution of cLD values in the 3D Hi-C dataset are displayed in Supplementary Figure S3.1. One can see that the average cLD of the int group is significantly higher than that in the no interaction regions. The corresponding numerical values supporting this figure are listed in Supplementary Table 3.2. The rigorous test to justify the significance statistically will be elaborated in section 3.4. This result indicates that cLD statistic captures the 3D interaction between genes. It is also clearly observed that the cLD decreases along with the increased distance between genes. In addition, the largest difference in cLD between int groups and no-int groups occurs in the first distance unit and the difference in cLD between int groups and no-int groups decreases considerably as the distance increases. This phenomenon implies that the interaction between gene pairs with shorter distances is more likely to result in a higher cLD value.

Then, in order to show that cLD is more appropriate than LD in capturing the 3D interaction between genes, we generated Supplementary Figure S3.2, the results analogue to Supplementary Figure S3.1 using the standard LD ($r^2$). The details of how we conducted the calculations are presented as follows. Again the gene pairs are split into the same 13 distance units. Then for each gene pair within the Hi-C assessed 3D-interaction, we randomly chose 2000 SNV pairs from it to calculate the average LD value. For each selected SNV pair, we calculated its distance and then among the gene pairs without Hi-C interaction we randomly

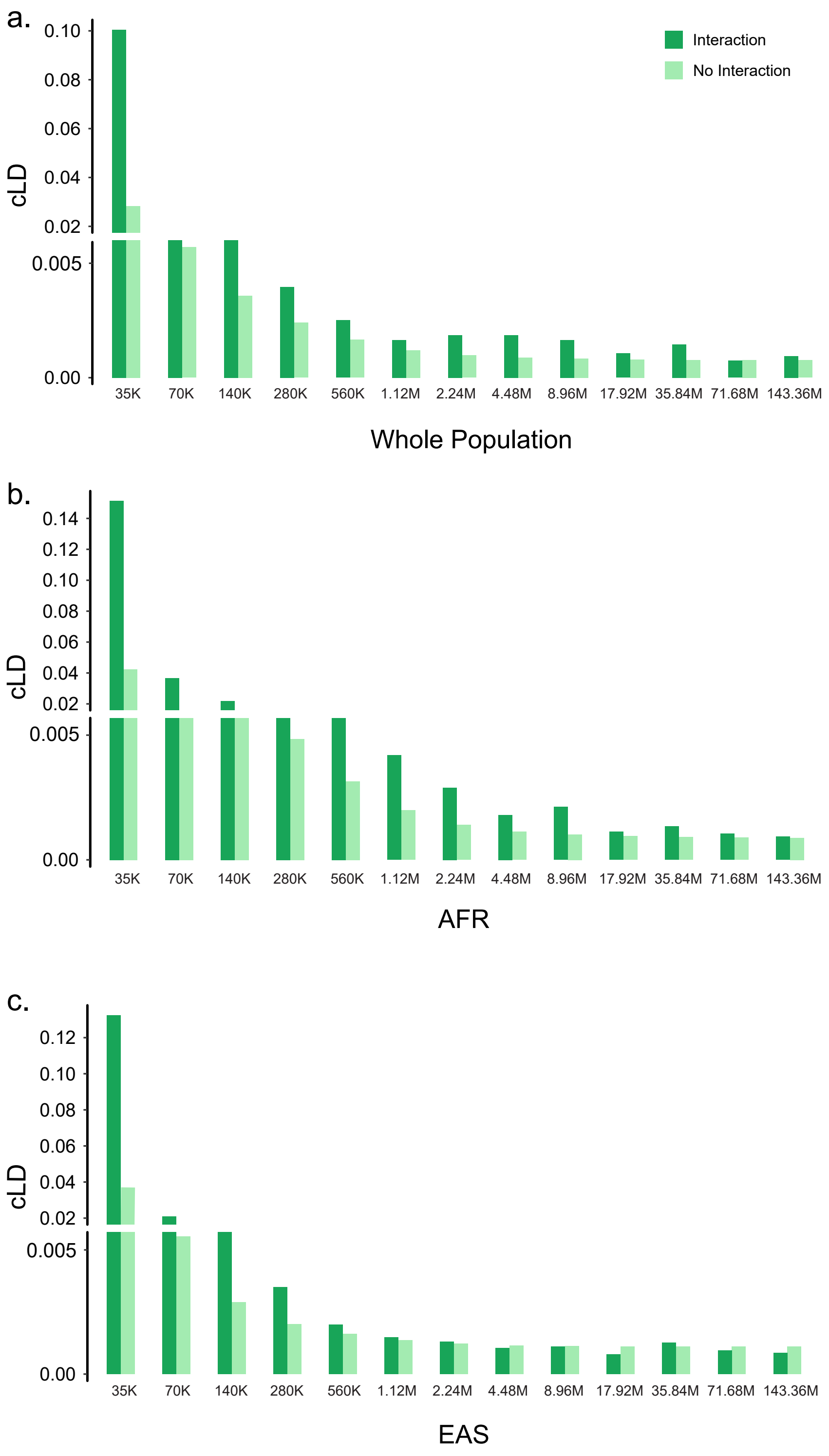

**Supplementary Fig. S3.1:** The comparisons of cLD values between the 3D chromatin interaction regions and non-interaction regions among 13 different distance groups in (a) the whole population, (b) AFR, and (c) EAS. The European population, EUR, has been displayed in the main text Figure 3(a)

selected another SNV pair with the same or very similar distance. As a result, we had total 2000 randomly selected SNVs from gene pairs without interaction that are paired up with the 2000 SNVs from gene pairs within 3D-interaction regions. Finally, we grouped the 2000 distance pairs into the 13 distance units and calculated the average LD for in interaction regions and non-interaction regions within each distance unit.

As shown in Supplementary Figure S3.2, the average LD values for the 3D-interaction and non-interaction regions are only slightly different in each of the 13 distance units. Indeed, statistically, the subtle difference is not significant, quantified by our two-sample test in section 3.4.

Therefore, based on the observations from Supplementary Figure S3.1 and Supplementary Figure S3.2, as well as their statistical test (to be presented in section 3.4), we concluded that cLD is a more appropriate statistic than LD to capture the 3D interaction between genes.

The 95% confidence intervals of the cLD in 3D chromatin interaction regions is :

From this Table S3.1, we can see the cLD's distribution of the interaction group is higher than the distribution of the no-interaction group. This means our cLD value does distinguish the interaction between genes.

#### 3.3 LD and cLD in databases for gene-gene interactions

In this section, we first gave an introduction to the gene-gene interaction databases used for assessing the enrichment of cLD in interacting gene pairs. Based on these interaction databases, the gene pairs are separated into interaction groups and no-interaction groups. Then we presented the results of mean cLD values in these two groups.

**1. Biogrid database**

Biogrid [4] is a biomedical interaction repository with data compiled through comprehen-

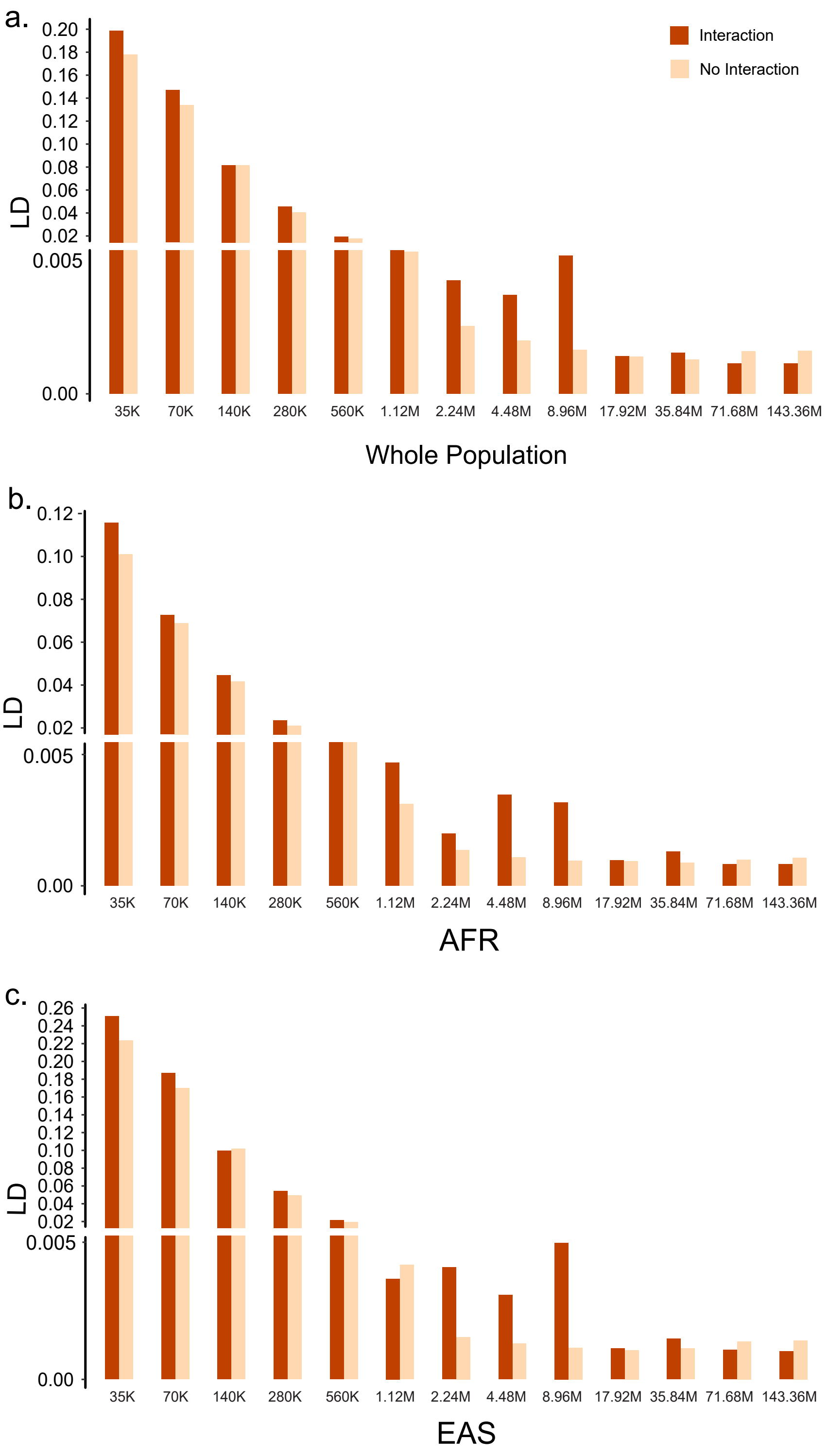

**Supplementary Fig. S3.2:** The comparisons of LD values between the 3D chromatin interaction regions and non-interaction regions among 13 different distance groups in (a) the whole population, (b) AFR, and (c) EAS. The European population, EUR, has been displayed in the main text Figure 3(b).

sive curation efforts. Their current index is version 4.4.197 and searches 76,687 publications for 2,045,743 protein and genetic interactions, 29,093 chemical interactions and 1,070,825 post-translational modifications from major model organism species. The data were downloaded from https://thebiogrid.org/.

**2. Reactom database**

Reactome[5] is a free, open-source, curated and peer-reviewed pathway database. This dataset provides intuitive bioinformatics tools for the visualization, interpretation and analysis of pathway knowledge to support basic research, genome analysis, modelling, systems biology and education. The data were downloaded from https://reactome.org/.

**3. MINT database**

MINT [6] focuses on experimentally verified protein-protein interactions mined from the scientific literature by expert curators.

Protein interaction databases represent unique tools to store, in a computer-readable form, the protein interaction information disseminated in the scientific literature. Well-organized and easily accessible databases permit the easy retrieval and analysis of large interaction data sets. MINT is a database designed to store data on functional interactions between proteins. Beyond cataloguing binary complexes, MINT was conceived to store other types of functional interactions, including enzymatic modifications of one of the partners. Since the MINT dataset provides interaction between proteins, we used the Uniprot website to convert the protein's UniProtKB to the gene's Ensembl ID. The data were downloaded from https://mint.bio.uniroma2.it/.

**4. Intact database**

The Intact database [7] is similar to the MINT dataset. Intact provides a freely available, open-source database system and analysis tools for molecular interaction data. All interactions are derived from literature curation or direct user submissions and are freely available. We first transferred the protein-protein interaction to gene-gene interaction and then conducted further analysis.

We combined the four interaction databases (Biogrid, Reactom, MINT and Intact) into a single data entry. Then we used the cLD statistic calculated before to analyze their distributions in the gene pairs in or not in the interaction databases. The results are displayed in Supplementary Figure S3.3, from which we observed that the cLD of gene pairs with interactions is higher than that of gene pairs without interaction. The statistical significance tests in section 3.4 also quantitatively confirmed this intuition. Thus we concluded that the cLD captures the gene-gene interaction well.

#### 3.4 Two-sample tests assessing significance levels

In this section, we compared the mean cLD (or LD) between the two categories under comparison, i.e., the interaction or non-interaction regions (in terms of either 3D interaction or gene-gene interactions) in a statistical rigorous way. More specifically, we designed the Mantel-Haenszel test and Fisher's exact test to assess the p-values to quantify the significant level.

The null hypothesis of the tests is that the distributions (of cLD, for example) are the same in the two groups (of gene pairs in interaction or non-interaction regions); and the alternative hypothesis is that the distributions are not the same. To distinguish these two hypotheses, we designed a 2 x 13 contingency table, in which 2 columns are for two distributions and 13 rows are for 13 distance groups. Each cell of the contingency table will be filled in the following way:

First we create a "standard" by calculating the median (i.e., the 0.5 quantile) value in the whole population (e.g., EUR or AFR), denoted by $q_{0.5}$. Then we define the "success rate" for a given distance group and a given distribution (interaction or non-interaction) as the proportion of statistics (such as cLD or LD) that are larger than the standard value of $q_{0.5}$. Then we put these 13 x 2 success rates into the contingency table.

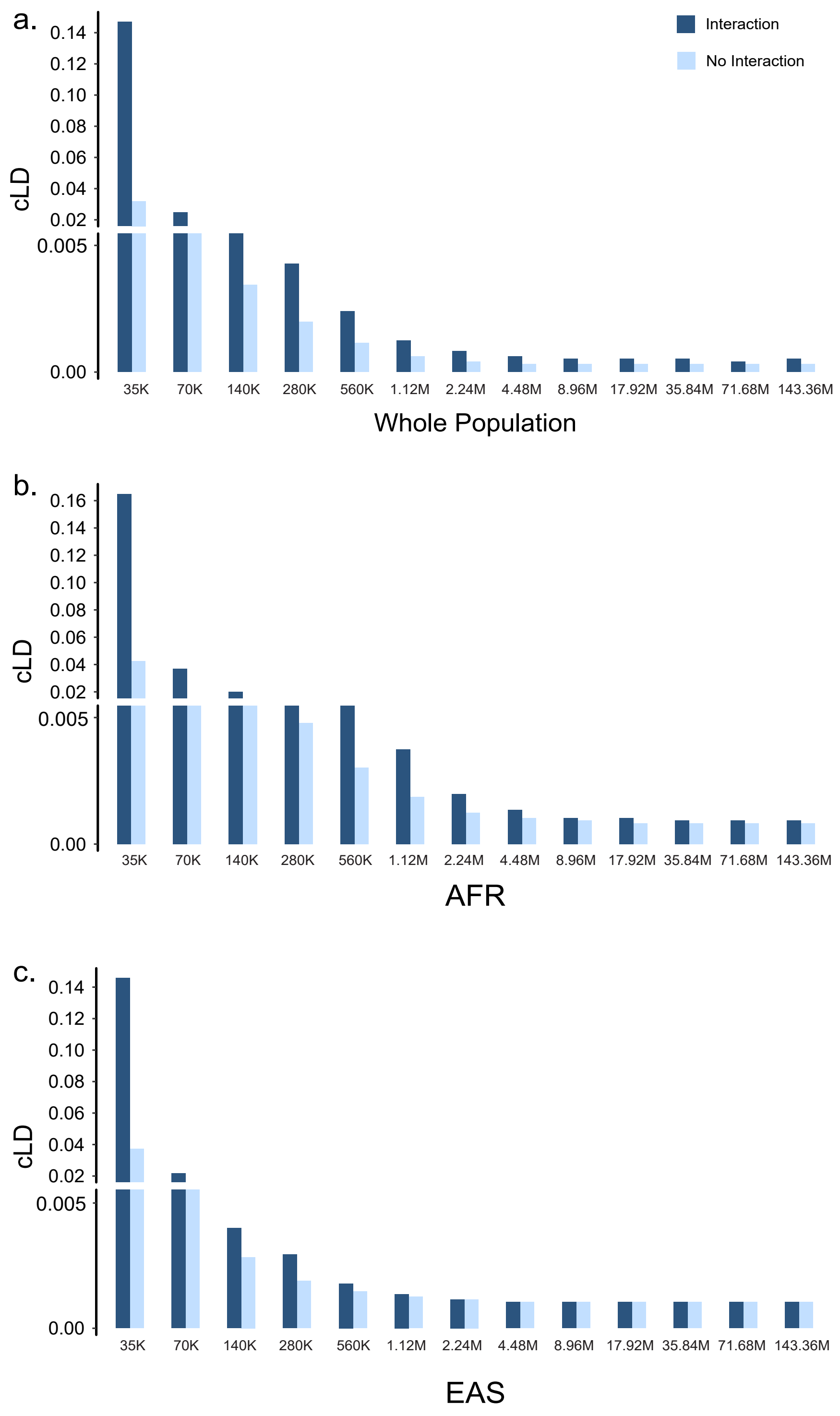

**Supplementary Fig. S3.3:** The comparisons of cLD values between the gene-gene interaction regions and regions without interactions among 13 different distance groups in (a) the whole population, (b) AFR, and (c) EAS. The European population, EUR, has been displayed in the main text Figure 4.

If we view the data collected from 13 different distance groups as independent experiments, then the standard Mantel-Haenszel test can be used to test the significance level. Meantime, Fisher's Exact test can also be utilized to test whether the two columns are from the same distribution.

By applying the above Mantel-Haenszel test and Fisher's Exact test to the cLD and LD distributions, we assessed whether this visible or subtle differences displayed in the figures in section 3.2 and section 3.3 are statistically significant.

##### 3.4.1 Two-sample Tests of cLD & LD in 3D interacting regions

We conducted the Mantel-Haenszel test and Fisher's exact test for LD and cLD in the Hi-C assessed 3D interaction regions. Since the results for different populations are quite similar, we only presented those for EUR in Table S3.2 as an illustration. The results for AFR, EAS and the whole population are listed in Supplementary Table 3.6 (enclosed the attached Excel file).

Based the protocol described in the previous subsection, we formed the data for the test in Supplementary Table S3.2 for EUR population. The last two columns (Int-ration and Non-int-ratio) are the contingency table ready for the two-sample test.

The Fisher's Exact test statistic is 29.84 and the Mantel-Haenszel test statistic is 12.99. The $p$-values for both tests are less than 1E-50 (based on the corresponding R libraries) and thus we rejected the null hypothesis based on either test. The Fisher's Exact and the Mantel-Haenszel test statistics are 51.34 and 28.52 respectively for EAS population, and 87.80 and 34.81 for AFR population. The $p$-values of these test statistics are also less than 1E-50. Therefore we concluded that, the cLD values in the 3D interaction regions are significantly higher than that of the non-interaction regions.

Similarly, for LD, again using EUR as the representative example, the number of successes, number of failures and success/failure ratios for interaction and non-interaction groups are presented in Supplementary Table S3.3.

Based on Supplementary Table S3.3 for the EUR population, the Fisher's Exact test statistic is $-45.65$ and the Mantel-Haenszel test statistic is $-45.68$. The $p$-values for both tests are very close to 1 and thus we could not reject the null hypothesis based on either test. The Fisher's Exact and the Mantel-Haenszel test statistics are $-21.34$ and $-21.35$ respectively for EAS population, and $-20.74$ and $-20.75$ for AFR population. The $p$-values of these test statistics are also very close to 1. Therefore we concluded that, the LD in the 3D-interaction regions are not higher than that of the non-interaction group. In other words, the high-LD gene pairs are not overlapped with the gene pairs within the 3D interaction regions, hence the standard LD could not capture the 3D interactions between gene pairs.

##### 3.4.2 Two-sample tests of cLD in gene-gene interaction databases

Analogue to the case of 3D-interactions, we conducted the same two-sample tests to the categorization based on gene-gene interactions. We combined all the reported interactions by the four datasets Biogrid, Reactom, MINT and Intact to form the interaction category and the compliment as the non-interaction category. The number of successes, number of failures and the success/failure ratios for interaction and non-interaction groups for EUR, AFR and EAS are presented in Supplementary Table 3.5 (enclosed the attached Excel file).

Based on the contingency table for the EUR population, the Fisher's Exact test statistic is 60.09 and the Mantel-Haenszel test statistic is 40.59. The $p$-values for both tests are less than 1E-50 and thus we rejected the null hypothesis based on either test. Based on the table for AFR population, the Fisher's Exact test statistic is 58.73 and the Mantel-Haenszel test statistic is 39.72. The $p$-values for both tests are again less than 1E-50 and thus we rejected the null hypothesis based on either test. Based on the table for EAS population, the Fisher's Exact test statistic is 87.80 and the Mantel-Haenszel test statistic is 34.81. The $p$-values for both tests are less than 1E-50 and thus we rejected the null hypothesis based on either test. Taking together, the above results proved that cLD captures the gene-gene interaction well.

#### 3.5 Enrichment of high-cLD pairs in 3D Interaction regions

In this section we provide details of the analysis of the enrichment of high-cLD pairs as a function of the extreme value of cLD, which is presented in the Main Text Figure 3c,d,e.

First, we set up multiple cLD cutoffs to distinguish different levels of extremely high values, i.e., 0.05, 0.1, 0.2, 0.3, and 0.4. Then, for the interval between every two consecutive cutoffs, we count gene pairs with their values falling in the interval and split them into regions covered by 3D interactions or not, respectively. As the number of gene pairs is huge, we only considered gene pairs on chromosome 1 and chromosome 2, which in contains 15,102,208 and 6,053,460 gene pairs, respectively. Amount these gene pairs, 1,686 and with 1,357 are within the 3D regions. Next, we calculated the percentage of gene pairs within 3D-interaction regions with the cLD values falling in the internal ($P_1$) as well as the percentage of such high-cLD gene pairs in the whole chromosome ($P_2$). The ratio between the above two percentages is plotted as the "Improved ratio" for this particular interval defined by two cLD-value cutoffs. The specific math formula are:

$$P_1 = \frac{Number\ of\ high\ cLD\ gene\ pairs\ within\ 3D\ regions}{Total\ number\ of\ gene\ pairs\ within\ 3D\ regions}$$

$$P_2 = \frac{Number\ of\ high\ cLD\ gene\ pairs\ in\ the\ chromosome}{Total\ number\ of\ gene\ pairs\ in\ the\ chromosome}$$

$$Improved\ ratio = \frac{P_1}{P_2}$$

Evidently, the improved ratios at different cLD cutoffs, across different populations (EUR, EAS and AFR), are large, and enjoy an increasing trend as the cLD cutoff increases (Main text Figure 3c,d,e; Supplementary Tables 3.7).

**Supplementary Table S3.1:** The confidence intervals of cLD values in the 3D chromatin interaction regions and non-interaction regions among 13 different distance groups in three populations.

| **EUR** | | | | | | |
|---|---|---|---|---|---|---|
| Distance groups | Int 0.025 | Int 0.5 | Int 0.975 | No Int 0.025 | No Int 0.5 | No Int 0.975 |
| 35K | 2e-5 | 0.0150 | 0.8712 | 2e-6 | 0.0017 | 0.4499 |
| 70K | 1e-5 | 0.0043 | 0.2516 | 3e-6 | 0.0007 | 0.0514 |
| 140K | 5e-6 | 0.0030 | 0.0543 | 3e-6 | 0.0005 | 0.0280 |
| 280K | 5e-6 | 0.0021 | 0.0276 | 2e-6 | 0.0004 | 0.0200 |
| 560K | 3e-6 | 0.0013 | 0.0182 | 2e-6 | 0.0003 | 0.0143 |
| 1.12M | 3e-6 | 0.0007 | 0.0136 | 2e-6 | 0.0003 | 0.0105 |
| 2.24M | 1e-6 | 0.0005 | 0.0081 | 9e-7 | 0.0003 | 0.0080 |
| 4.48M | 1e-6 | 0.0005 | 0.0060 | 7e-7 | 0.0002 | 0.0063 |
| 8.96M | 2e-6 | 0.0004 | 0.0126 | 6e-7 | 0.0002 | 0.0060 |
| 17.92M | 3e-6 | 0.0004 | 0.0051 | 5e-7 | 0.0002 | 0.0061 |
| 35.84M | 1e-6 | 0.0004 | 0.0078 | 5e-7 | 0.0002 | 0.0059 |
| 71.88M | 1e-6 | 0.0004 | 0.0067 | 5e-7 | 0.0002 | 0.0060 |
| 143.36M | 1e-6 | 0.0006 | 0.0064 | 6e-7 | 0.0002 | 0.0057 |
| **AFR** | | | | | | |
| Distance groups | Int 0.025 | Int 0.5 | Int 0.975 | No Int 0.025 | No Int 0.5 | No Int 0.975 |
| 35K | 5e-5 | 0.0238 | 0.9068 | 2e-6 | 0.0039 | 0.4356 |
| 70K | 7e-5 | 0.0119 | 0.3047 | 2e-6 | 0.0020 | 0.0657 |
| 140K | 3e-5 | 0.0083 | 0.0783 | 2e-6 | 0.0012 | 0.0454 |
| 280K | 7e-6 | 0.0049 | 0.0461 | 2e-6 | 0.0007 | 0.0313 |
| 560K | 1e-5 | 0.0034 | 0.0300 | 2e-6 | 0.0005 | 0.0202 |
| 1.12M | 3e-6 | 0.0016 | 0.0192 | 2e-6 | 0.0004 | 0.0120 |
| 2.24M | 1e-6 | 0.0008 | 0.0103 | 1e-6 | 0.0003 | 0.0080 |
| 4.48M | 1e-6 | 0.0005 | 0.0070 | 1e-6 | 0.0003 | 0.0061 |
| 8.96M | 1e-6 | 0.0004 | 0.0056 | 9e-7 | 0.0003 | 0.0054 |
| 17.92M | 1e-6 | 0.0004 | 0.0054 | 8e-7 | 0.0002 | 0.0049 |
| 35.84M | 8e-7 | 0.0004 | 0.0048 | 9e-7 | 0.0002 | 0.0047 |
| 71.88M | 8e-7 | 0.0004 | 0.0046 | 8e-7 | 0.0002 | 0.0046 |
| 143.36M | 7e-7 | 0.0004 | 0.0044 | 6e-7 | 0.0002 | 0.0045 |
| **EAS** | | | | | | |
| Distance groups | Int 0.025 | Int 0.5 | Int 0.975 | No Int 0.025 | No Int 0.5 | No Int 0.975 |
| 35K | 1e-5 | 0.0106 | 0.8201 | 2e-6 | 0.0007 | 0.4352 |
| 70K | 7e-6 | 0.0017 | 0.2281 | 2e-6 | 0.0004 | 0.0324 |
| 140K | 4e-6 | 0.0013 | 0.0402 | 1e-6 | 0.0004 | 0.0162 |
| 280K | 3e-6 | 0.0009 | 0.0166 | 1e-6 | 0.0003 | 0.0117 |
| 560K | 2e-6 | 0.0007 | 0.0098 | 1e-6 | 0.0002 | 0.0092 |
| 1.12M | 2e-6 | 0.0005 | 0.0074 | 5e-7 | 0.0003 | 0.0074 |
| 2.24M | 1e-6 | 0.0005 | 0.0057 | 4e-7 | 0.0003 | 0.0065 |
| 4.48M | 9e-7 | 0.0003 | 0.0056 | 4e-7 | 0.0003 | 0.0060 |
| 8.96M | 1e-6 | 0.0004 | 0.0053 | 3e-7 | 0.0002 | 0.0059 |
| 17.92M | 1e-6 | 0.0003 | 0.0046 | 4e-7 | 0.0002 | 0.0058 |
| 35.84M | 2e-6 | 0.0004 | 0.0047 | 3e-7 | 0.0002 | 0.0057 |
| 71.88M | 1e-6 | 0.0004 | 0.0043 | 3e-7 | 0.0002 | 0.0057 |
| 143.36M | 1e-7 | 0.0004 | 0.0036 | 3e-7 | 0.0002 | 0.0057 |

**Supplementary Table S3.2:** EUR counts for Mantel-Haenszel test (3D-interaction, cLD)

| | Int-succ | Int-fail | Non-int-succ | Non-int-fail | Int-ratio | Non-int-ratio |
|---|---|---|---|---|---|---|
| 35K | 75 | 7 | 47642 | 21004 | .91 | .69 |
| 70K | 209 | 12 | 45516 | 19804 | .95 | .70 |
| 140K | 318 | 44 | 82967 | 38725 | .88 | .68 |
| 280K | 379 | 51 | 148833 | 78943 | .88 | .65 |
| 560K | 337 | 68 | 260934 | 163980 | .83 | .61 |
| 1.12M | 100 | 38 | 451231 | 335797 | .72 | .57 |
| 2.24M | 51 | 31 | 779888 | 661248 | .62 | .54 |
| 4.48M | 43 | 20 | 1346386 | 1238435 | .68 | .52 |
| 8.96M | 41 | 30 | 2389097 | 2281671 | .57 | .51 |
| 17.92M | 72 | 46 | 4126180 | 4073582 | .61 | .50 |
| 35.84M | 122 | 65 | 6792988 | 6878739 | .65 | .49 |
| 71.88M | 154 | 54 | 9759922 | 10001248 | .68 | .49 |
| 143.36M | 158 | 72 | 11206696 | 11646643 | .53 | .49 |
| Overall | 2059 | 538 | 37438280 | 37439819 | .79 | .50 |

**Supplementary Table S3.3:** EUR counts for Mantel-Haenszel test (Hi-C, LD)

| | Int-succ | Int-fail | Non-int-succ | Noint-fail | Int-ratio | Non-int-ratio |
|---|---|---|---|---|---|---|
| 35K | 764 | 43 | 807 | 0 | .94 | 1. |
| 70K | 1786 | 365 | 2151 | 0 | .83 | 1. |
| 140K | 2854 | 1480 | 4323 | 11 | .65 | 1. |
| 280K | 2137 | 3024 | 4515 | 646 | .41 | .87 |
| 560K | 720 | 3506 | 1420 | 2808 | .17 | .34 |
| 1.12M | 123 | 2019 | 9 | 2133 | .06 | 0 |
| 2.24M | 41 | 956 | 0 | 999 | .02 | 0 |
| 4.48M | 15 | 556 | 0 | 572 | .02 | 0 |
| 8.96M | 26 | 403 | 0 | 430 | .06 | 01 |
| 17.92M | 0 | 234 | 0 | 234 | 0 | 0 |
| 35.84M | 0 | 260 | 0 | 261 | 0 | 0 |
| 71.88M | 0 | 258 | 0 | 117 | 0 | 0 |
| 143.36M | 0 | 117 | 0 | 117 | 0 | 0 |
| Overall | 8466 | 13221 | 13225 | 8462 | .39 | .61 |

### Chapter 4

### Protein interactions and docking

Despite the large number of gene-gene interactions reported by the four databases used in section 3.3, many of the gene pairs with high cLD are still not found in these databases, such as between 3BCZ and 4RIQ (encoded for by genes MEMO1 and DPY30, respectively) with a cLD of 0.86. This leaves room for discovering novel gene-gene interactions indicated by high cLD values. In this chapter, we presented our characterization of the novel interactions by computational prediction of protein docking. This enables us to investigate the nature of the relationship between the two cLD-identified genes through the receptor-ligand interactions of their translated proteins.

#### 4.1 Pipeline for protein-protein docking analysis

The interacting protein-protein candidates were selected by identifying the top gene pairs with cLD values between the ranges of 0.33 to 0.86. Subsequently, the proteins' tertiary structures were obtained from the RCSB Protein Data Bank (PDB) [8]. The structures were validated based on their resolution being less than 3Å, having almost zero Ramachandran outliers, and the macro-molecules parent organism being Homo sapiens. The structures were visualized and prepared for docking using PyMOL [9].

We used Huang Lab's software HDOCKlite for protein-protein docking. HDOCKlite is a user downloadable software-based version of their more famous, CASP-CAPRI winning HDOCK server. HDOCK's docking protocol uses a hybrid protein docking technique that incorporates both ab-initio and homology-based principles [10, 11]. We selected HDOCKlite as its a time-tested and validated user-friendly tool, and it can run on High-Performance Computing (HPC) clusters. Additionally, it had very short processing times enabling us to conduct multiple analyses efficiently. The docking analysis was conducted using HDOCK-lite's default parameters.

Once the docking analysis was conducted between the candidate pairs, post-processing of the protein-protein complexes was conducted. First, it was ensured that the formed complexes had negative binding energies. Based on the laws of thermodynamics, the more negative the value of the binding energy, the more stable the protein substrate complex formed between the two proteins [10]. Afterwards, the 3D complex was examined in PyMOL and the interacting surfaces of the two macro-molecules were investigated using DIMPLOT from LigPlot+ [12]. DIMPLOT provides a 2D representation of the interacting surfaces at an atomic level complete with the chemical bonds responsible for the establishment of the protein-protein complex.

#### 4.2 Protein-protein docking results

We conducted protein docking analysis for the 19 pairs of genes with large cLD values (top 0.01% among all gene pairs) with cMAF $> 0.05$ and existing IDs in PDB, however, not reported in any databases Supplementary Table S4.1. As examples, we selected five pairs of protein-protein docking results for presentation. The detailed list of the candidate proteins and their cLD values as well as binding energy are listed below in Supplementary Table S4.2. The graphical overviews of the formed complexes are depicted in this section as Supplementary Figures S4.1, S4.2, S4.3 and S4.4 except that of the 3BCZ and 4RIQ complex which is present in the Main Text Figure 5.

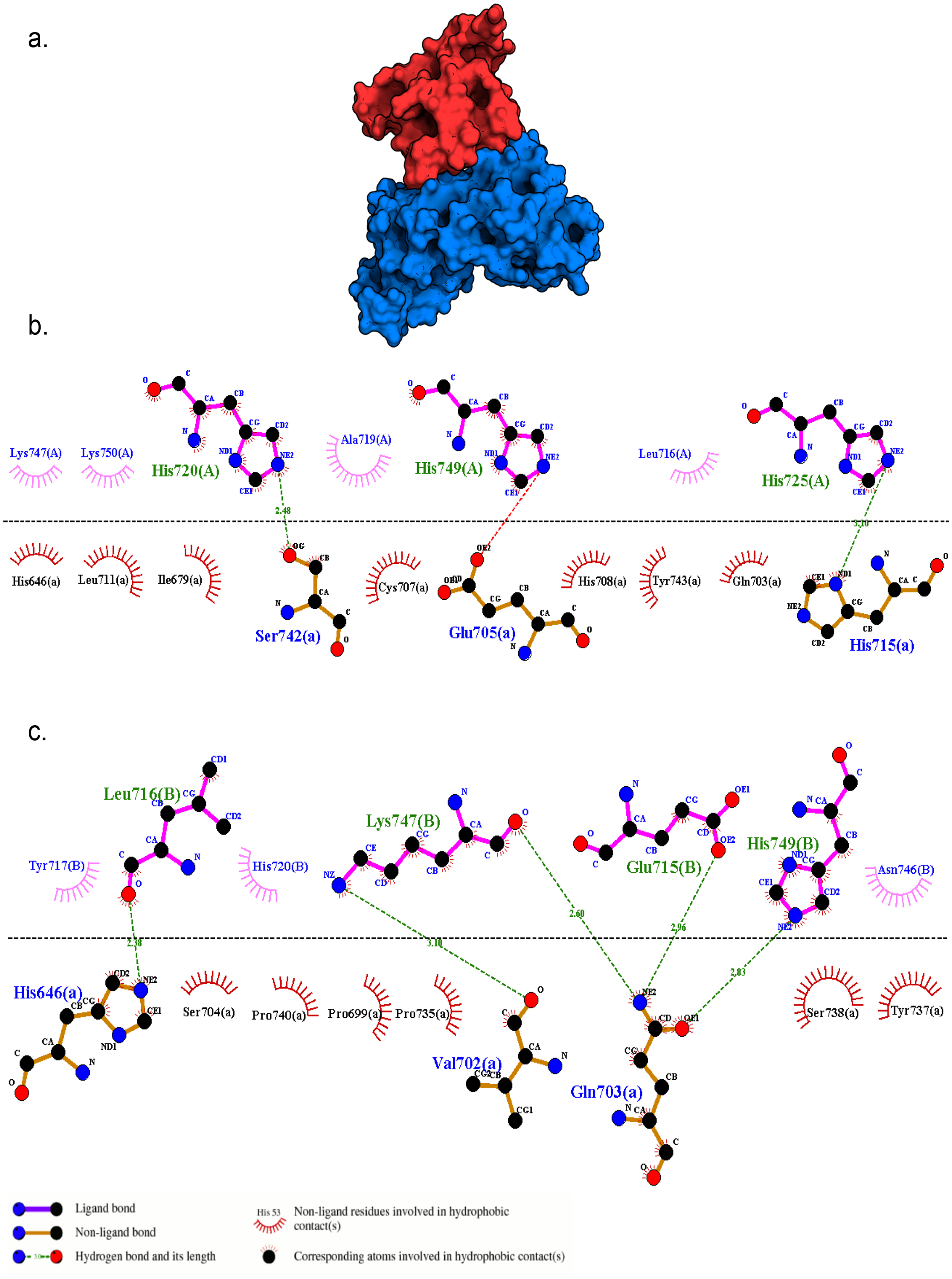

**Supplementary Fig. S4.1:** Protein docking interaction between 1FYV and 4OM revealed by cLD (0.69) with a binding affinity of -266.77 kJ/mol. a) Structure of 1FYV (red) and 4OM (blue) protein-protein complex. b-c) 2D representation of closest interacting residues around the protein-protein interaction interfaces, including hydrogen bonds (green dotted line) and hydrophobic interactions (read and rose semi-circle with spikes). Residues for the 1FYV are depicted by uppercase letters (A, B) and for the 4OM are depicted by lower lowercase letter (a).

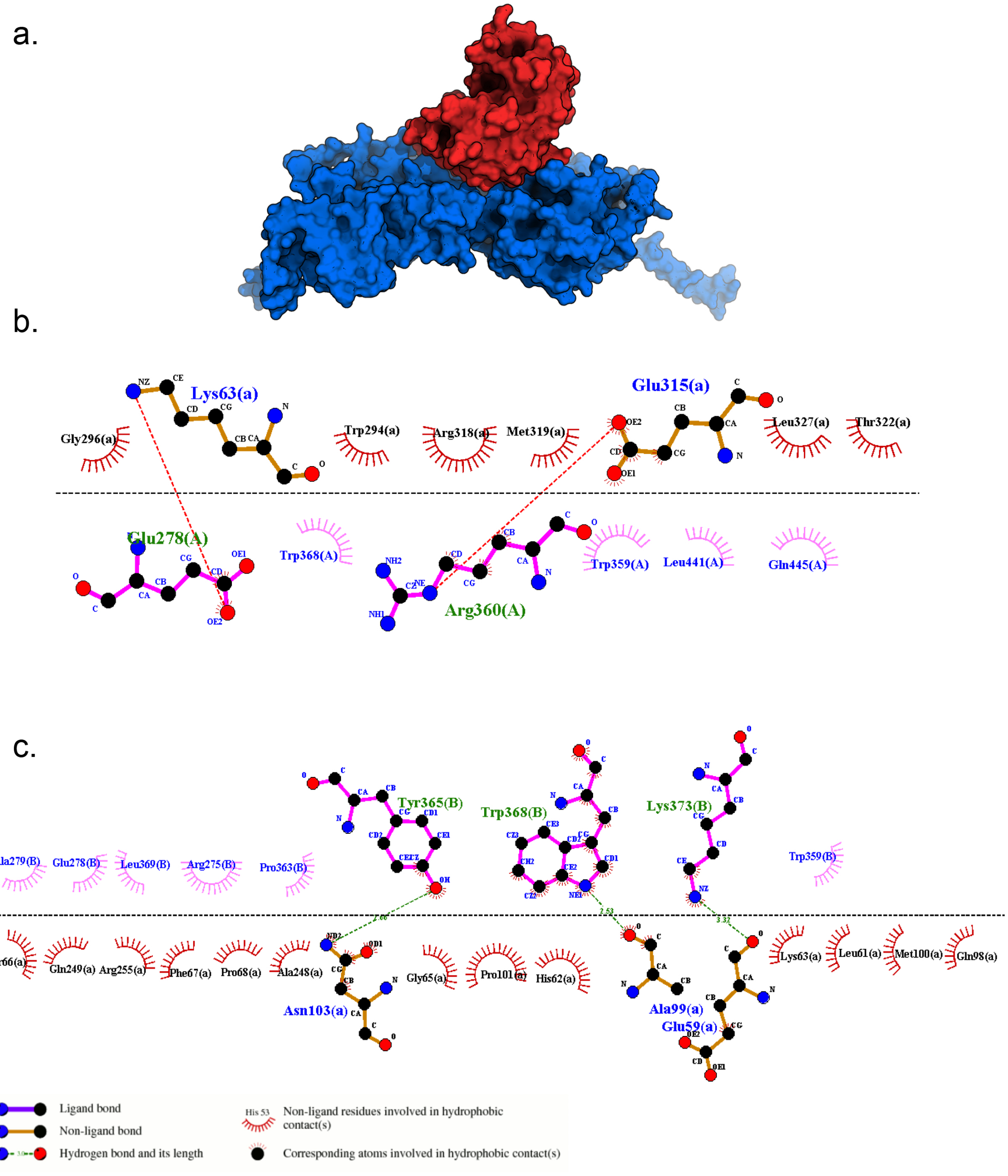

**Supplementary Fig. S4.2:** Protein docking interaction between 3S5N and 4HND revealed by cLD (0.52) with a binding affinity of -302.64 kJ/mol. a) Structure of 3S5N (red) and 4HND (blue) protein-protein complex. b-c) 2D representation of closest interacting residues around the protein-protein interaction interfaces, including hydrogen bonds (green dotted line) and hydrophobic interactions (read and rose semi-circle with spikes). Residues for the 3S5N are depicted by uppercase letter (A) and for the 4HND are depicted by lowercase letters (a, b).

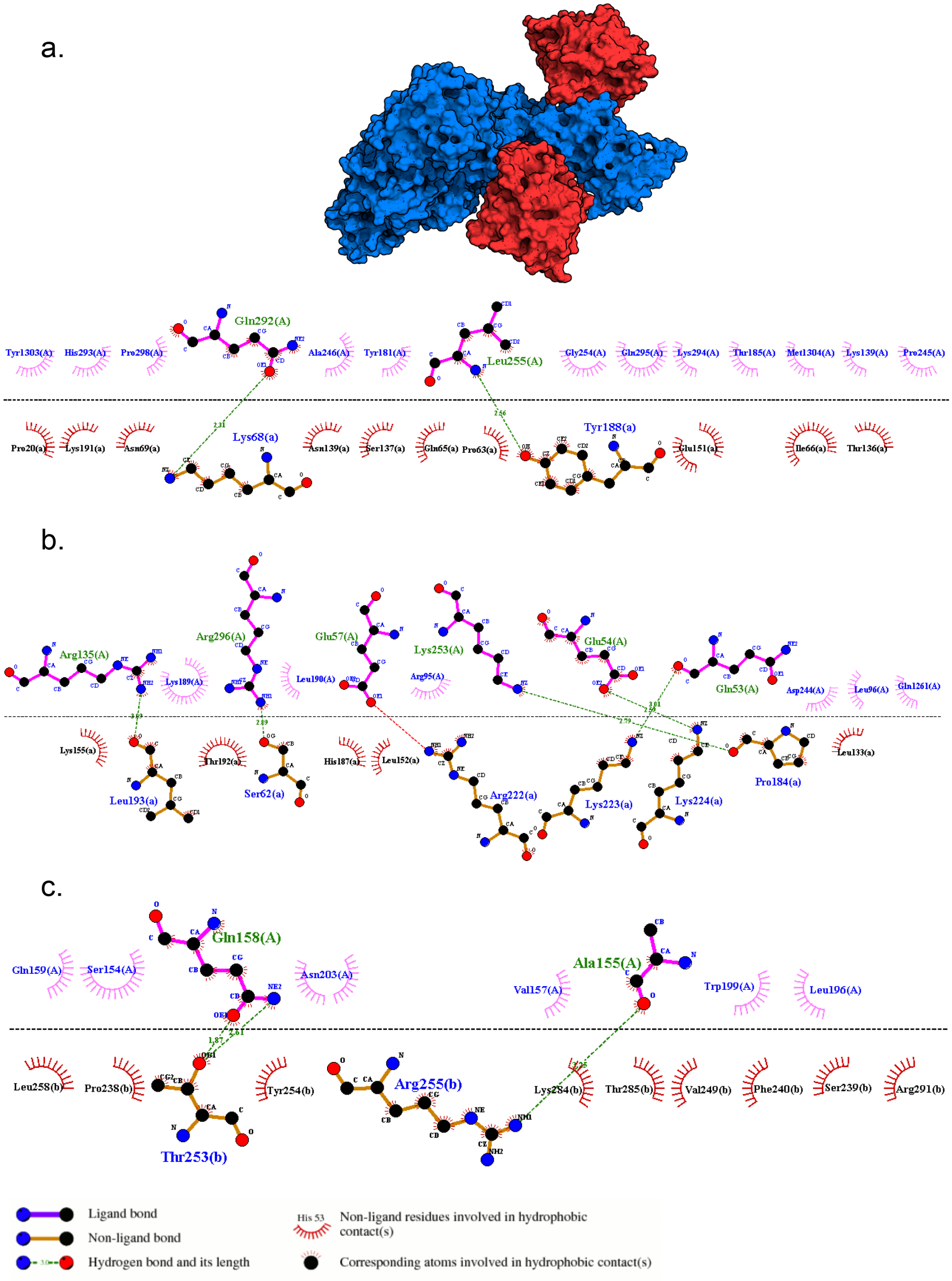

**Supplementary Fig. S4.3:** Protein docking interaction between 1WVA and 6HO2 revealed by cLD (0.40) with a binding affinity of -263.19 kJ/mol. a) Structure of 1WVA (red) and 6HO2 (blue) protein-protein complex. b-c) 2D representation of closest interacting residues around the protein-protein interaction interfaces, including hydrogen bonds (green dotted line) and hydrophobic interactions (read and rose semi-circle with spikes). Residues for the 1WVA are depicted by uppercase letter (I) and for the 6HO2 are depicted in lowercase letter (a).

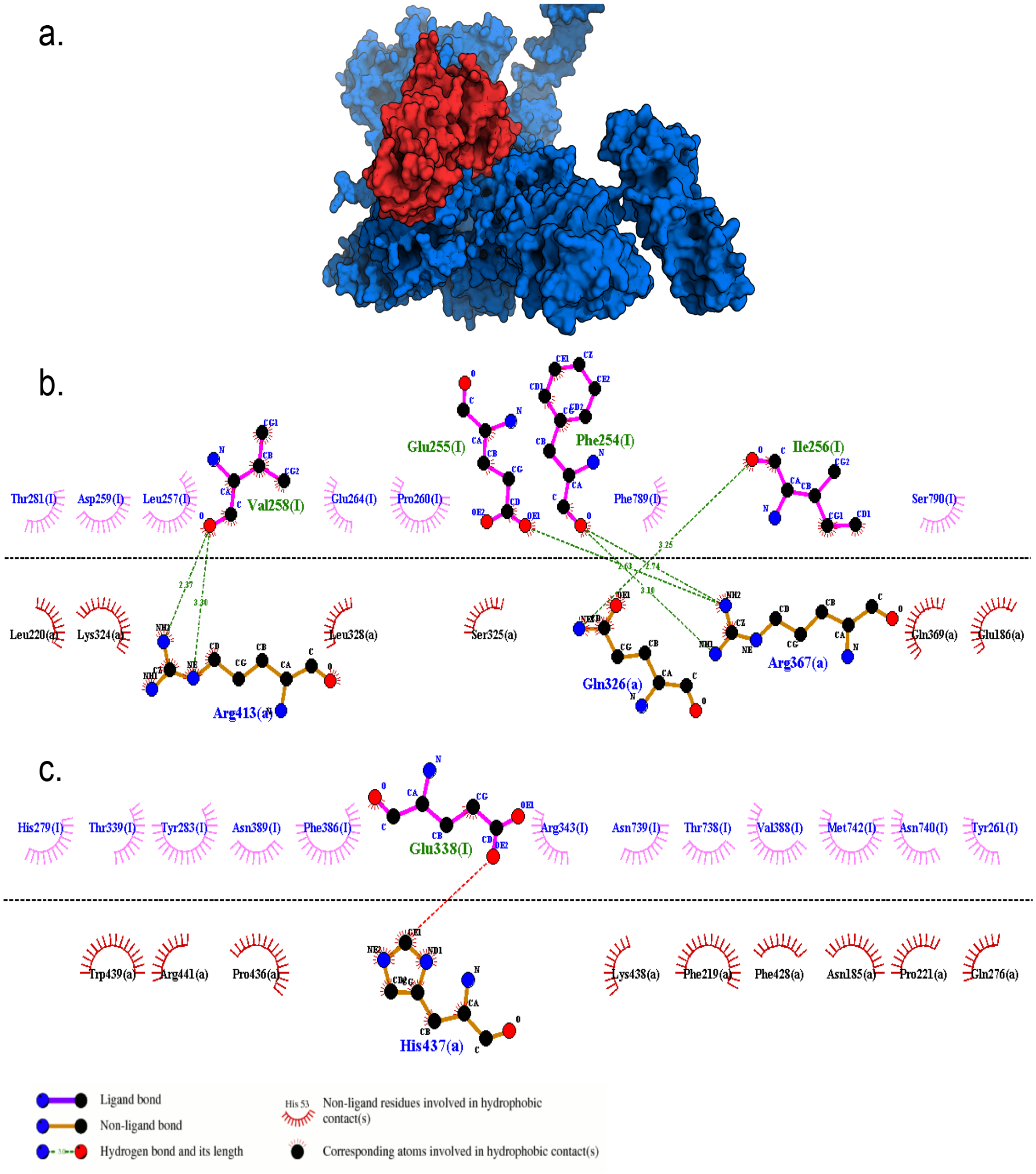

**Supplementary Fig. S4.4:** Protein docking interaction between 3L81 and 5FUR revealed by cLD (0.33) with a binding affinity of -277.36 kJ/mol. a) Structure of 3L81 (red) and 5FUR (blue) protein-protein complex. b-c) 2D representation of closest interacting residues around the protein-protein interaction interfaces, including hydrogen bonds (green dotted line) and hydrophobic interactions (read and rose semi-circle with spikes). Residues for the 3L81 are depicted by uppercase letters (A, B) and for the 5FUR are depicted in the lowercase letter (a).

**Supplementary Table S4.1:** List of 19 gene pairs(not reported in any databases) with large cLD values with cMAF > 0.05 and existing IDs in PDB.

| Ensembl ID 1 | PDB ID 1 | Ensembl ID 2 | PDB ID 2 |
|---|---|---|---|
| ENSG00000162959 | 3BCZ | ENSG00000162961 | 4RIQ |
| ENSG00000114779 | 1IMJ | ENSG00000243989 | 1Q7L |
| ENSG00000168827 | 6VLZ | ENSG00000079257 | 2BO9 |
| ENSG00000174125 | 1FYV | ENSG00000174130 | 4OM7 |
| ENSG00000131187 | 6SZW | ENSG00000198055 | 2ACX |
| ENSG00000166278 | 2I6Q | ENSG00000243649 | 1DLE |
| ENSG00000240065 | 6AVO | ENSG00000168394 | 1JJ7 |
| ENSG00000231389 | 3LQZ | ENSG00000223865 | 3LQZ |
| ENSG00000124596 | 2EEE | ENSG00000001167 | 4AWL |
| ENSG00000118520 | 1WVA | ENSG00000112282 | 6H02 |
| ENSG00000241685 | 6YW7 | ENSG00000130429 | 6UHC |
| ENSG00000221838 | 3L81 | ENSG00000106290 | 5FUR |
| ENSG00000087077 | 1X61 | ENSG00000087085 | 1B41 |
| ENSG00000197448 | 1YZX | ENSG00000106144 | 6GKF |
| ENSG00000133742 | 1AZM | ENSG00000164879 | 1Z93 |
| ENSG00000042832 | 6SCJ | ENSG00000155926 | 2CUD |
| ENSG00000122705 | 6E5N | ENSG00000159921 | 2YHW |
| ENSG00000095380 | 1WVO | ENSG00000106785 | 6JBM |
| ENSG00000241935 | 3S5N | ENSG00000155252 | 4HND |

**Supplementary Table S4.2:** List of candidate proteins with their respective cLD values and binding affinities. All candidates formed stable protein-protein complexes with negative binding energies.

| Ensembl ID | Gene Name | PDB ID | Ensembl ID | Gene Name | PDB ID |
|---|---|---|---|---|---|
| ENSG00000162959 | *MEMO1* | 3BCZ | ENSG00000162961 | *DPY30* | 4RIQ |
| ENSG00000174125 | *TLR1* | 1FYV | ENSG00000174130 | *TRL6* | 4OM7 |
| ENSG00000241935 | *HOGA*1 | 3S5N | ENSG00000155252 | *PI4K2A* | 4HND |
| ENSG00000118520 | *ARG1* | 1WVA | ENSG00000112282 | *MED23* | 6H02 |
| ENSG00000221838 | *AP4M1* | 3L81 | ENSG00000106290 | *TAF6* | 5FUR |

### Chapter 5

### Association mapping using cLD and annotations

In this chapter, we tested whether cLD serves as a valid tool to discover genes whose interactions may play critical role in diseases. More specifically, we conducted analysis to assess whether gene pairs with highly differential cLD($\Delta$cLD) values distinguishing cases and controls are enriched in functionally relevant databases as well as sensible gene ontology (GO) and pathways (reported by the KEGG repository).

#### 5.1 Data source and quality control

We calculated cLD on Autism Spectrum Disorder (ASD) whole exon sequencing dataset [18], which is downloaded from dbGaP web portal (http://www.ncbi.nlm.nih.gov, ID: phs000298.v4.p3). The dataset is quality controlled using PLINK [19]. The QC steps include excluding markers with missing rate $> 0.1$, high deviations from Hardy-Weinberg Equilibrium $> 10\text{-}6$, and removing samples with missing rate $> 0.1$.

#### 5.2 Calculating cLD in case-control datasets

We conducted the analysis composed of the following 5 steps:

Step 1: For the target whole exome sequencing dataset, we only retained rare variants with MAF $< 0.005$

Step 2: We then located gene region for all human genes, and the corresponding gene information is downloaded from GENCODE (https://www.gencodegenes.org/human/ ). The gene region is ranging from location start to location end.

Step 3: For the target dataset, if there is no rare variant within the gene region, this gene will be excluded, otherwise, we further split the entire dataset into case and control groups. We then calculated the cMAF within each gene region for cases and controls separately.

Step 4: We retained genes with cMAF $> 0.05$ in both case and control groups.

Step 5: Lastly, we calculated cLD within the gene regions for each gene pair. This calculation was also done for case and control separately.

#### 5.3 Enrichment of cLD-differential genes in functional databases

We calculated the absolute differences between the cLD values in cases and controls for every pairs of genes. This difference is the contrast between cases and controls in terms of cLD indicated interactions. We then ranked them from largest to smallest. From the ranked list, we selected the top 200, 500, 1,000, 1,500, or 2,000 gene pairs for functional annotations. For each set of high $\Delta$cLD pairs, we form a gene list containing all genes included by at least a pair.

The model for the functional annotation is to check the enrichment of cLD identified genes in known databases. We referred to two databases: one is from Simons Foundation Autism Research Initiative (SFARI) [20] and the other one is from DisGeNet [21]. The SFARI database is a well-established repository for existing ASD genes [20] and its data can be directly downloaded. For the DisGeNet database, we used "Autism" as the keyword and searched for all Autism-associated genes. We checked these two databases separately. For a given protocol, we defined the "success" as the number of identified genes reported in the corresponding database, and the "success rate" as the ratio of these validated genes against all of the selected genes.

To test whether the enrichment is significant. We calculated the $p$-values under hypergeometric distribution. The density of the hypergeometric distribution with parameters $m$, $n$, $k$, and $N = m + n$, is given by the formula below:

$$P(x) = \frac{\binom{m}{x}\binom{n}{k-x}}{\binom{N}{k}}$$

$m$: The number gene is the database. The database is downloaded from either SFARI or DisGeNet.

$n$: The total number of genes in the population ($= N$) minus the number of genes in the database ($= m$). Here, $N$ =20,000, as we considered roughly 20,000 human genes in the population.

$k$: The number of genes from selected top (200, 500, ..., 2000) gene pairs.

$q$: The number of selected genes being validated in the database.

We calculated the distribution function using `"phyper"`, provided in R. The $p$-value is calculated as `1- phyper(q, m, n, k, lower.tail = TRUE, log.p = FALSE)`. If the $p$-value $< 0.05$, it is considered as a significant enrichment.

By applying the above hypergeometric test to the previously formed $\Delta$cLD gene lists and the DisGeNet and SFARI databases, we could quantify the enrichment led by different sets of high $\Delta$cLD genes. It is observed that, as we included more top gene pairs (from 200 to 2,000), the number of success is increasing in both DisGeNet and SFARI databases. The success rate is ranging from 9.80% to 10.34% for the DisGeNet database, the success rate increased a little bit as we included more genes. For the SFARI database, the success rate is ranging from 6.72% to 8.84%, which is a bit lower than the success rate in the DisGeNet database. This might be due to the reason that firmly verified ASD genes in the SFARI database are limited (Supplementary Table 5.2). Looking at the enrichment p-value in both databases, it is evident that the p-values are significant for all different selection of gene pairs (from top 200 to top 2,000), and the p-value is decreasing (i.e., more significant) dramatically as we include more gene pairs (Main Text Figure 6a, 6b and Supplementary Table 5.2). This indicated that high $\Delta$cLD genes between the case and control may be highly associated with diseases.

From the top 10 gene pairs with the highest cLD values, we identified 20 unique genes. 14 out of these 20 genes (70%) have been reported to be associated with ASD, including *DENND4A*, *EFCAB5*, *ABI2*, *RAPH1*, *MSTO1*, *DAP3*, *ARL13B*, *PRB2*, *PRB1*, *ZNF276*, *FANCA*, *ADAM7*, *SLC26A1* and *TUBB8* (Supplementary Table S5.1). Among the rest of six genes, some genes also showed indirect associations with ASD. For example, mutations of *RAB11A* were identified in patients with developmental diseases or/and epilepsy [13]. Gene *IDUA* participates in the degradation of glycosaminoglycans (GAGs) within the lysosome [14], and degradation of glycosaminoglycans has been suggested to be occurred in ASD patients [15]. Long noncoding RNA has been widely reported to be associated with ASD [16, 17], therefore *RP11-1148O4.2* , *RP11-624C23.1* and *DHRS4-AS1* may consider as potential candidate genes for ASD.

**Supplementary Table S5.1:** From the top 10 gene pairs with the highest cLD values, we identified 20 unique genes. 14 out of these 20 genes (70%) have been reported to be associated with ASD, including *DENND4A, EFCAB5, ABI2, RAPH1, MSTO1, DAP3, ARL13B, PRB2, PRB1, ZNF276, FANCA, ADAM7, SLC26A1* and *TUBB8*.

| Gene 1 | Gene 2 | cLD | Reference 1 | Reference 2 |
|---|---|---|---|---|
| RAB11A | DENND4A | 0.223 | | PMID: 30559488 |
| EFCAB5 | RP11-1148O4.2 | 0.200 | PMID: 26189338<br>PMID: 22542183<br>PMID: 24267886 | |
| ABI2 | RAPH1 | 0.188 | PMID: 30610205<br>PMID: 15542242<br>PMID: 33157009 | PMID: 26731442 |
| MSTO1 | DAP3 | 0.170 | PMID: 31463572<br>PMID: 28544275 | PMID: 28392909<br>PMID: 25363768 |
| ARL13B | DHFRL1 | 0.165 | PMID: 23153492 | |
| PRB2 | PRB1 | 0.159 | PMID: 31540669<br>PMID: 27708715 | PMID: 31540669<br>PMID: 27708715 |
| ZNF276 | FANCA | 0.157 | PMID: 31838722<br>PMID: 33431980 | PMID: 25531569 |
| ADAM7 | RP11-624C23.1 | 0.146 | PMID: 33374371 | |
| IDUA | SLC26A1 | 0.141 | | PMID: 33374967<br>PMID: 26399424 |
| TUBB8 | DHRS4-AS1 | 0.134 | PMID: 22169095 | |

#### 5.4 Enrichment of cLD-differential genes in GO and KEGG

As another independent line of analysis in parallel to the DisGeNet and SFARI databases, we conducted gene ontology, or GO, enrichment [22] and KEGG pathway analysis [23]. The GO enrichment is carried out from the aspect of biological process (BP) [22]. KEGG is used to systematically analyze gene functions, which links gene information with high-level functional information [24]. The GO enrichment and KEGG pathway analyses were conducted using the 'clusterProfiler' R package [25]. The enriched GO terms and KEGG pathways with a $p$-value $< 0.05$ were considered disease-related biologic processes or signalling pathways.

We included all selected genes within the top 2000 gene pairs with high $\Delta$cLD for the GO enrichment and KEGG pathway analysis. We showed that genes with high $\Delta$cLD are enriched in sensible pathways and biological processes that are related to ASD (Main Text Figure 6c, 6d). Here are examples supported by literature:

Glutamatergic synapse was the leading pathway identified in KEGG analysis (Main Text Figure 6c). Glutamate is the major excitatory neurotransmitter in the central nervous system and its dysfunction has been central to neurotransmitters that are involved in autism[26]. A fruitful of GWAS studies have reported the associations between genes involved in glutamatergic neurotransmission and ASD [27–29]. Moreover, glutamatergic synapses play a key role in neuronal functions, including synaptic transmission, neuronal migration, excitability, plasticity, long-term potentiation, and long-term depression [30]. Many genes, including SHANK families [31–33], NLGN and NRXN families [34] related to the glutamatergic synapses have been found to be associated with ASD [35].

Followed by the Glutamatergic synapse, the second most enriched pathway is the focal adhesion. Many neuronal cell adhesion molecules help form the structure of the neuronal network during development and are involved in cognitive functions and memory [36]. For ASD, a body of studies have also demonstrated that alterations in synaptic genes including those encoding cell adhesion molecules, e.g., cadherins, NCAM, and neurexin superfamily, can affect the structural connectivity of neurons, thus play important roles in the pathogenesis of ASD [37–39].

From the most significantly enriched GO terms, we observed extracellular matrix organization, extracellular structure organization and cell-substrate adhesion are associated with the structural integrity of the neuronal network. Other biological processes, including dendrite development, dendrite morphogenesis and synapse organization, are all contributing to synaptic dysfunction, neurotransmission, and neurodevelopment (Main Text Figure 6d).
